# Investigating the effect of acute consumption of kiwifruit on cognitive function using functional MRI

**DOI:** 10.64898/2026.11.09.765279

**Authors:** Michael Asghar, Sally Eldeghaidy, Simone Poggesi, Joanne Hort, Susan Francis

## Abstract

**Objectives:** Acute nutritional interventions may modulate cognitive function and underlying brain activity. This study investigated the impact of consuming three kiwifruit varieties—green (Zespri Green), gold (Zespri SunGold), and red (Zespri RubyRed)—on cognitive performance and brain activation using functional magnetic resonance imaging (fMRI).

**Methods:** Fourteen healthy participants completed a randomized, repeated-measures study involving four MRI sessions following consumption of each kiwifruit type and a maltodextrin control. Cognitive function was assessed using an N-back working memory task during fMRI acquisition. Behavioural performance, psychometric measures (mood, fatigue, appetite), and blood biomarkers (vitamin C) were also collected.

**Results:** Kiwifruit consumption significantly increased plasma vitamin C levels relative to control, with peak concentrations observed at 100 minutes post-consumption. Behaviourally, task difficulty increased with N-back load, while psychometric measures indicated greater positive affect following kiwifruit compared to control. fMRI results showed that the maltodextrin condition elicited broader brain activation, suggesting reduced neural efficiency. In contrast, kiwifruit— particularly the red variety—showed enhanced differentiation between low and high cognitive load (2-back vs. 0-back), with increased activation in regions associated with working memory and cognitive control.

**Conclusions:** These findings suggest that acute kiwifruit consumption may enhance cognitive efficiency and modulate brain responses, with effects varying by kiwifruit cultivar.

## Introduction

Over the last decade, a growing body of research has highlighted the broad health benefits of kiwifruit consumption, linking regular intake to improvements in nutritional status as well as psychological well-being, and digestive, immune, and metabolic health. Kiwifruit is rich in micronutrients, particularly vitamin C [1] and polyphenols [2], both of which are known to influence physiological and cognitive processes. Commercially, three main varieties of kiwifruit are available: green (Zespri Green), gold/yellow (Zespri SunGold), and the more recent red cultivar (Zespri RubyRed). These cultivars differ in their nutritional composition, with RubyRed containing the highest levels of Vitamin C (~189 mg/100g), followed by SunGold (~152mg/100g) and green (~88mg/100g), alongside elevated phenolic content.

Emerging evidence suggests that vitamin C status and kiwifruit consumption are associated with enhanced cognitive performance and mood. Higher plasma vitamin C levels have been linked to improved cognitive processing [3], while supplementation through kiwifruit intake has been shown to improve mood in young adults [4]. Specifically, SunGold kiwifruit has been shown to reduce mood disturbances in mood-compromised adults [5], restore vitamin C saturation in participants with respiratory infections, and reduce fatigue and depression symptoms [6]. Consumption of green kiwifruit has been associated with improved sleep quality, potentially due to its high level of folate and serotonin content [7].

Polyphenols, another key component of kiwifruit, have been associated with improved microvascular function [8] and reduced risk of neurodegeneration and cognitive decline [9]. Neuroimaging studies further demonstrate that polyphenol-rich interventions can modulate brain activity and enhance cognitive performance. For example, concord grape juice consumption has been associated with improved recognition memory and increased activation in prefrontal and parietal regions using functional Magnetic Reso-nance Imaging (fMRI) with Blood Oxygenation Level Dependent (BOLD) contrast [10]; while higher consumption of ulam (a traditional vegetable salad) has been linked to increased dorsolateral prefrontal cortex (DLPFC) activation during a working memory (N-back) task [11]. Similarly, consumption of whole coffee cherry extract has been shown to lead to deactivation in frontolimbic regions in a 7T fMRI study using a go-no-go task implying greater cognitive efficiency, while the placebo increased activity in regions dedicated to decision making [12].

In this study, we use fMRI-derived cortical metrics to investigate the neural correlates of cognitive processing following consumption of three kiwifruit varieties: green, gold (“SunGold”) and red (“RubyRed”). The primary aim was to assess the effect of consumption of kiwifruit on cognitive function using a working memory N-back letter task. We hypothesised that cognitive performance and associated brain activation would vary by kiwifruit variety based on their micronutrient content, with a graded effect of Red > Gold > Green > Maltodextrin control.

## Methods

### Participants and Study Design

Fourteen healthy participants (7 female; mean age ± SD: 28 ± 7 years) were recruited from the University of Nottingham via poster advertisements. Ethical approval was obtained from the Faculty of Medicine and Health Sciences Research Ethics Committee (FMHS2140223). All participants provided written informed consent and completed MRI safety screening and medical questionnaires prior to participation. Inclusion and exclusion criteria are detailed in the Supplementary Methods. The sample size of 14 participants used for this within-subject factorial fMRI design is in line with functional food neuroimaging paradigms using tightly controlled, compact cohorts [13–15].

Participants completed a familiarisation session, a sensory evaluation session, and four MR visits in a randomized, repeated-measures design. Each MRI visit included the randomised consumption of either a green (Zespri Green), gold (Zespri SunGold), red (Zespri RubyRed) kiwifruit, or a maltodextrin control matched for caloric content. All sessions were completed within a 51-day period (May–July).

### Familiarisation and Sensory Measures

As consumers in the UK are not overly familiar with gold and red varieties of Kiwi fruit a familiarisation session was performed such that any perception of novelty was already accounted for. During this session, participants were given two kiwifruits of each variety (Red, Gold, Green) for home consumption (Figure 1). In the sensory evaluation session, participants consumed one of each kiwifruit variety and completed questionnaires assessing overall liking [16] and emotional associations. The questionnaires rated perceptions of emotions using implicit testing lexicons developed at Massey University, NZ with Red, Gold and Green kiwifruit consumers.

**Figure 1:**
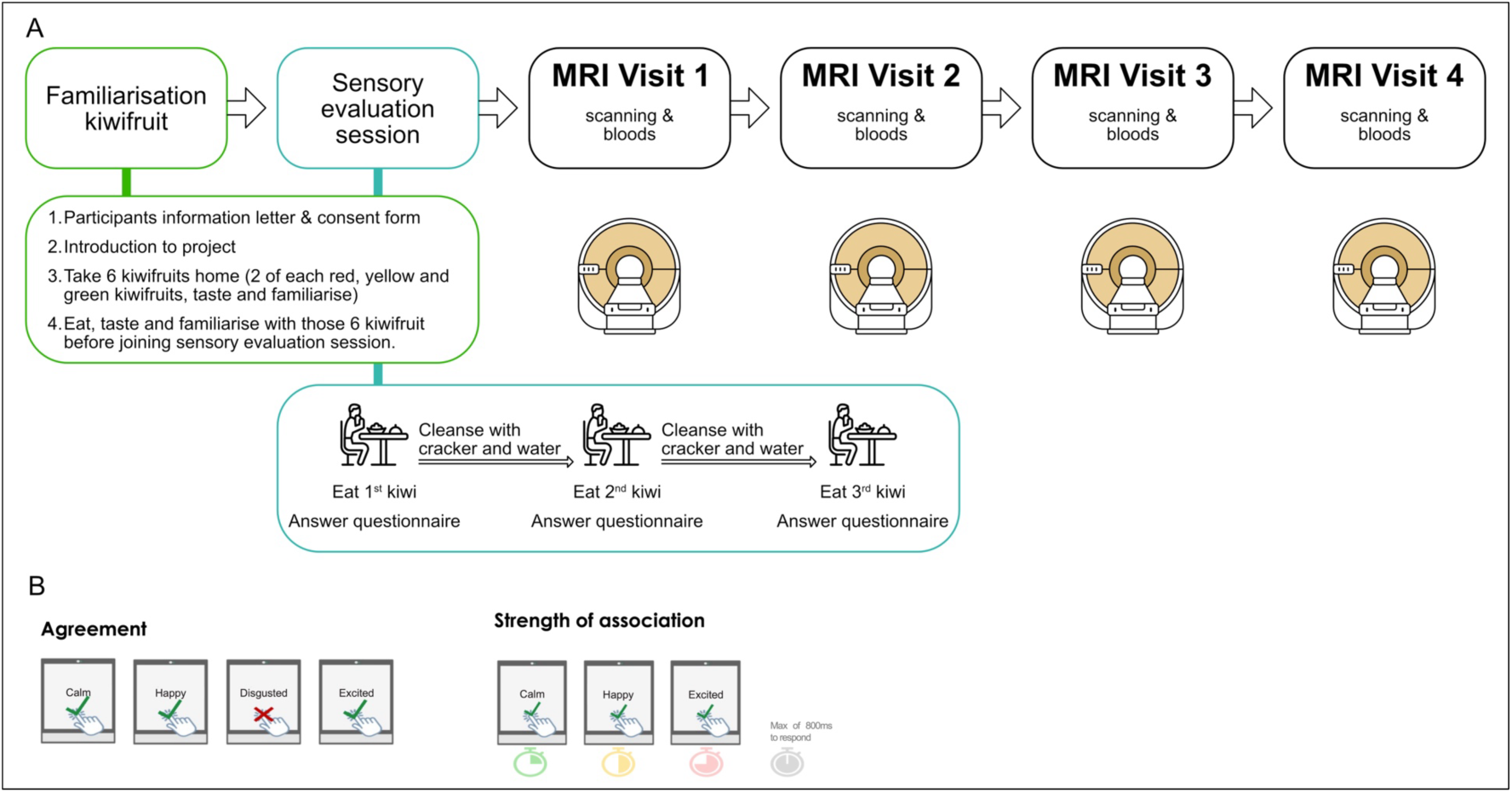
A) Familiarisation and sensory evaluation protocol, followed by four MRI visits. Kiwifruit (Gold, Green, Red) was randomised across MRI visit or a maltodextrin control matched for caloric content. B) Agreement and strength of association measures in implicit testing.

Participants indicated agreement with presented attributes via button press within 800 ms. The proportion of positive responses (“Go%”) represented agreement, while reaction time reflected strength of association (faster responses indicating stronger associations [17]). Non-responses were interpreted as weak or absent associations.

### MR Visit Protocol

Participants followed dietary restrictions for 24 hours prior to each MRI visit, avoiding foods high in vitamin C and polyphenols (e.g., fruits, vegetables, tea, coffee, chocolate, alcohol, nuts). Permitted foods included refined carbohydrates, meat, fish, and low-fat dairy. Water intake was unrestricted.

At each visit, instrumental measures of kiwifruit firmness and sweetness were taken. The firmness was measured using a non-destructive Firmness Tester (Willowbank Electronics Ltd, 2024) and sweetness was measured using a Brix device (in °Brix) to calculate sugar content; any kiwifruit outside of the normal acceptable range was discarded.

Participants consumed two kiwifruits at a timepoint immediately prior to scanning after having their baseline blood sampled. During the control visit, a maltodextrin drink matched for the same calorific content as the kiwifruit was consumed.

Psychometric assessments of the PANAS-20 (Positive and Negative Affect Schedule), MFSI-SF (Multidimensional Fatigue Symptom Inventory Short Form), and VAS (Visual Analog Scale) appetite scale were completed at three timepoints (pre-, mid-, and post-scan) during the MRI visit protocol as shown in Figure 2. Blood samples were collected before and after the scan to study vitamin C levels.

**Figure 2:**
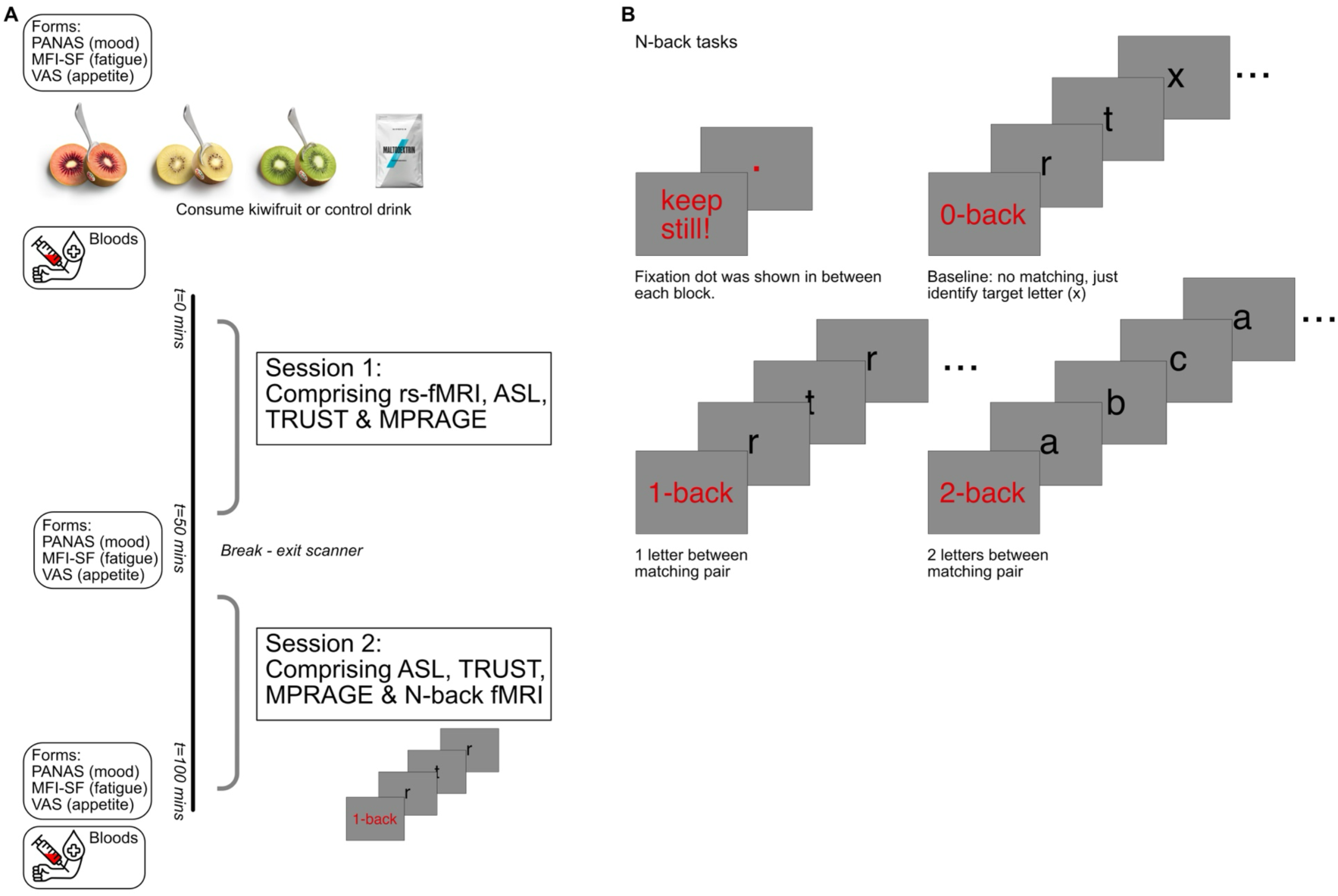
A) MRI visit protocol. rs-fMRI = resting state functional magnetic resonance imaging; N-back fMRI = N-back functional magnetic resonance imaging; ASL = arterial spin labelling (measure cerebral blood perfusion); TRUST = T2 relaxation under spin tagging (measure blood oxygen saturation fraction). B) N-back fMRI task shown illustrating the 0, 1, and 2-back conditions. Note: This paper focusses on results of the fMRI analyses. Participants also completed questionnaires and underwent venipuncture at fixed timepoints.

### MR Data Acquisition

MRI data were acquired on a 3T Philips Achieva scanner. Task-based functional imaging used BOLD EPI (TR = 2 s, voxel size = 3×3×3 mm^3^, duration ~10 min), alongside a structural MPRAGE scan for Freesurfer segmentation, and registration. Distortion correction (spin-echo EPI, blip-up/down) and dynamic noise scans were acquired for TOPUP correction [18] and thermal denoising using NOise Reduction with DIstribution Corrected (NORDIC) principal component analysis (PCA) [19]. Physiological signals (cardiac and respiratory) were recorded using a peripheral pulse unit (PPU) and pneumatic respiratory belt for physiological noise correction using RETROICOR [20]. Participants viewed stimuli via prism glasses and responded using a button box. Participants also underwent a resting state fMRI (rs-fMRI) scan comprising 20 minutes duration of 400 dynamics of double-echo gradient-echo echo-planar-imaging (GE-EPI), TE=25/45 ms, 3 mm isotropic spatial resolution, TR 1.5s.

Cognitive performance was assessed using a block-design N-back working memory task (0-, 1-, and 2-back conditions) (Figure 2). Participants were instructed to press a button whenever they saw a letter that was repeated with a gap of 1 letter (1-back), or 2 letters (2-back) between them. For the “0-back” task, the participant was instructed to press the button when they saw the letter “x”. At the start of each block, an instruction screen was presented for 2 s which displayed the forthcoming task condition. During a task period, 15 letters, 4 of which were targets, were presented for 1 s each in a pseudorandom order, with 1 s of blank screen between sequential letters. Each task period was followed by a 15 s rest period, during which a fixation dot was shown on the screen. The instruction screen, task period, and rest period formed an experimental block lasting (30+2+15) = 47 s. The experiment comprised of 4 blocks per condition (3 conditions of 0, 1, or 2-back) in each run = 564 s = 9.4 minutes.

### Data Analysis

#### Sensory evaluation and Implicit scores

Overall liking, Firmness and Brix (sugar content) were analysed using a one-way ANOVA and post-hoc pairwise comparisons were conducted using Tukey’s honestly significant difference (HSD) criterion to control for multiple comparisons. Implicit scores were derived from reaction time, and categorised into high (≥80th percentile), medium (50–80th percentile), and low (<50th percentile) associations.

#### fMRI processing

Task-fMRI data were pre-processed using RETROICOR for physiological noise correction, NORDIC PCA for thermal noise correction and FSL TOPUP for distortion correction. Quality assurance for fMRI data was assessed by calculating temporal SNR (tSNR) maps. N-back fMRI datasets were then analysed in Statistical Parametric Mapping 12 (SPM12 dev, UCL). Data with head motion > 1mm were excluded; 1 subject (4 visits) was removed due to excessive motion, and 9 other visits from other subjects, resulting in 43/56 usable imaging visits.

Analysis of the N-back data was performed in SPM12 using a general linear model (GLM) to derive group fMRI maps (across subjects) of the task condition (0-back, 1-back, 2-back) following acute consumption of gold, green, and red kiwifruit, and maltodextrin). Functional activation maps were parcellated using the Harvard-Oxford atlas. The main effect of interest was the 2-back versus. 0-back contrast to assesses the brain responses to the most versus least difficult task.

Behavioural performance (task accuracy calculated by computing the percentage correct and reaction time) was analysed across N-back conditions for each visit.

Resting state fMRI data were pre-processed in CONN toolbox (RRID:SCR_009550) [21] using the default processing pipeline comprising realignment, registration, normalization to MNI 2mm space. Denoising was performed using aCompCor [22], scrubbing and temporal band pass filtering [0.008 0.09Hz]. Group-ICA (independent component analysis) was performed in CONN using default parameters, (40 components, fast-ICA algorithm).

#### Blood analysis

Plasma vitamin C ascorbate levels were measured from blood samples at fasted and 100-minute timepoints.

## Results

### Sensory evaluation

There were no significant differences in overall liking (mean scores: Red 7.07, Green 7.29, Gold 7.70; Red vs Green p=0.91; Red vs Gold, p=1; Green vs Gold, p=0.91). There were no significant differences in brix scores (sugar content) (Red vs Green p=0.26; Red vs Gold, p=0.90; Green vs Gold, p=0.12) observed between the three kiwifruit varieties (one-way ANOVA; Figure 3). However, Green kiwifruits were firmer than Gold kiwifruits (p<0.007). Red kiwifruit did not differ significantly in firmness compared to Green (p=0.32) or Gold (p=0.20) kiwifruit.

**Figure 3:**
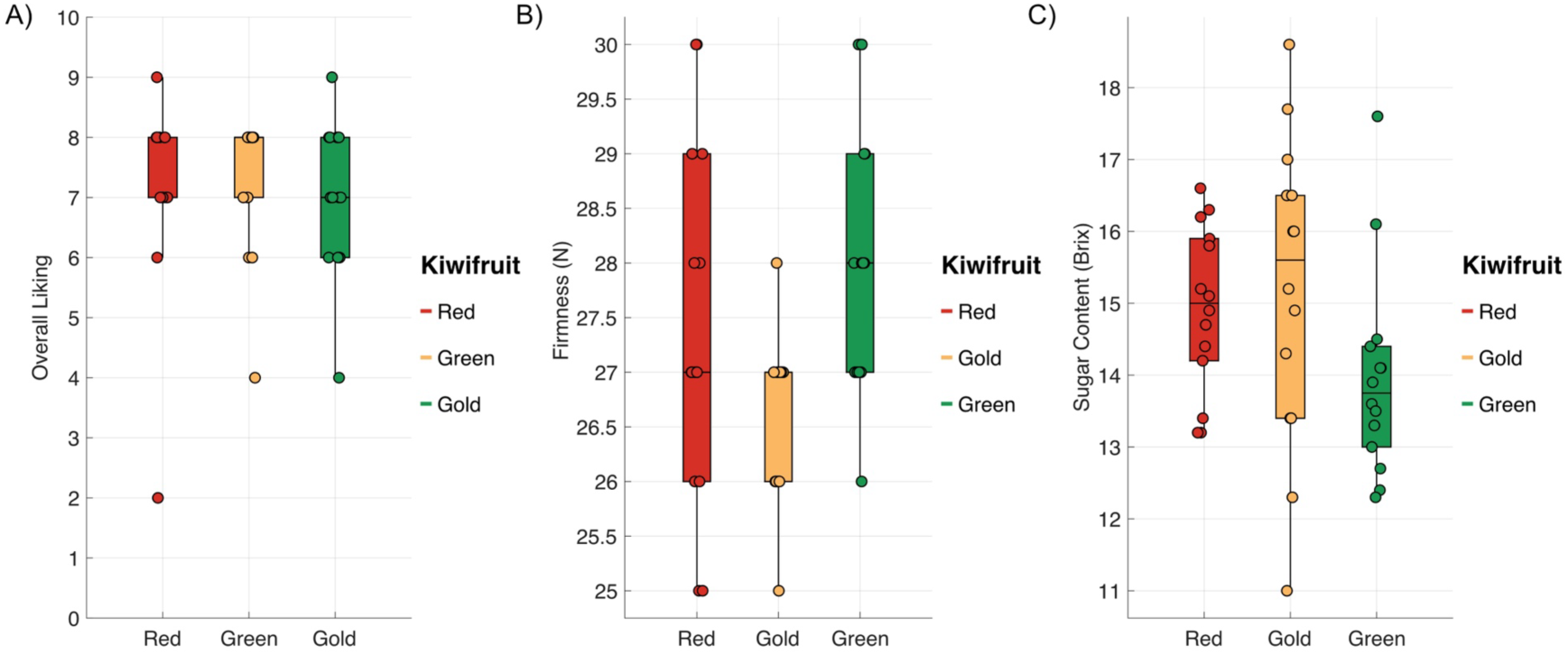
A) Overall liking, B) Firmness, and C) Brix scores for the kiwifruit varieties.

### Implicit association test

No negative emotional associations were reported for any sample. Implicit association testing revealed distinct positive emotional profiles: gold kiwifruit was primarily associated with ‘Satisfied’; green kiwifruit with ‘Happy’, ‘Pleasant’, and ‘Satisfied’; and red kiwifruit with ‘Excited’, ‘Happy’, and ‘Pleasant’.

### Questionnaires

PANAS-20 scores were significantly higher at the baseline (fasted) timepoint compared to mid and end timepoints. Additionally, scores were higher following consumption of red, green, and gold kiwifruit compared to the maltodextrin control. No significant effects of timepoint or visit were observed for MFSI-SF scores. For the VAS appetite score, the end timepoint had larger values than early and mid-timepoints. There was no difference across kiwifruit. There was no significant interaction between timepoint (start, mid, end) and intervention (red, gold, green, control). Results from the multi-way ANOVA (p < 0.05, corrected for multiple comparisons) are shown in Figure 4.

**Figure 4:**
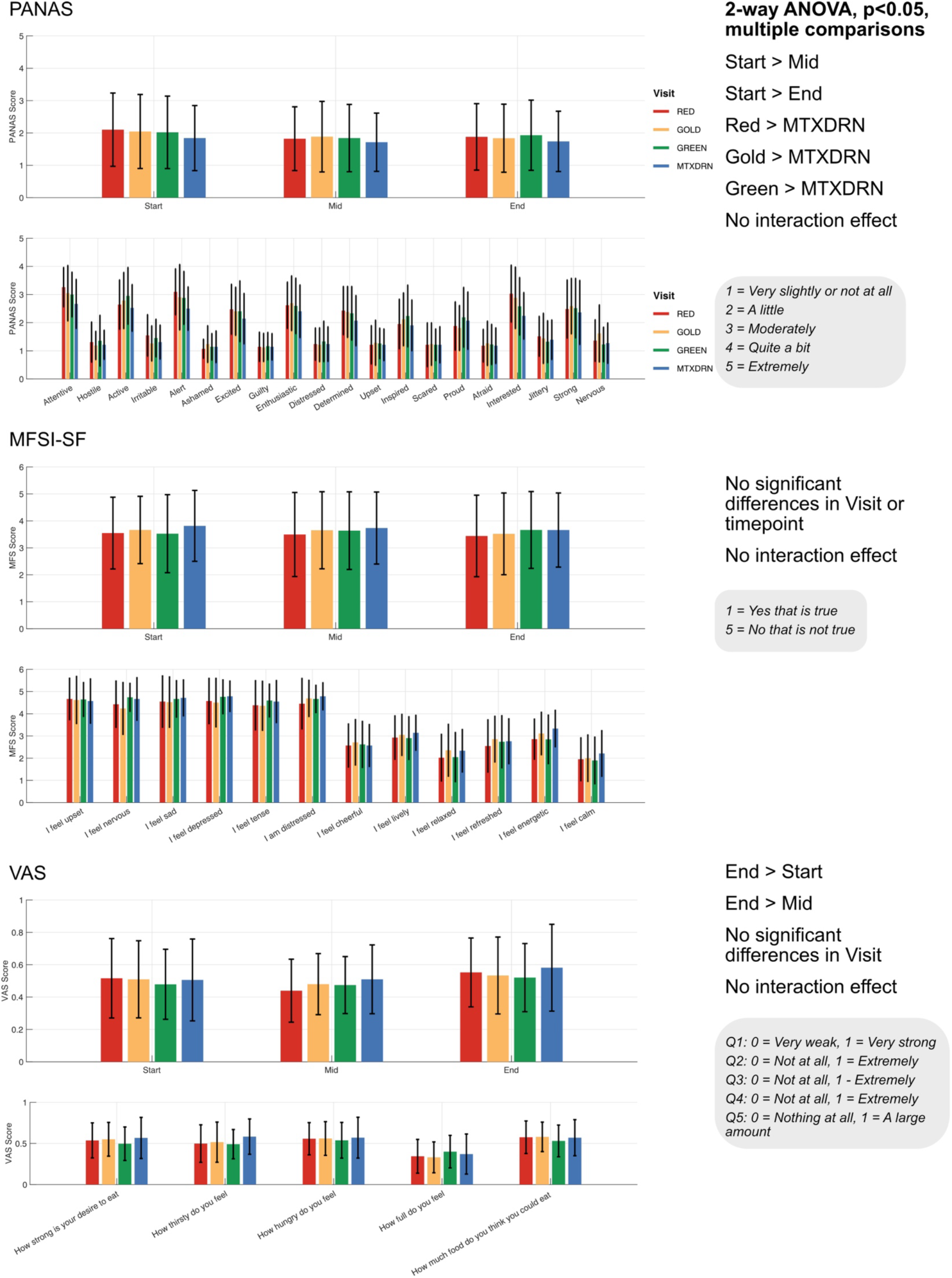
Results from A) Positive and Negative Affect Schedule (PANAS-20), B) Multidimensional Fatigue Symptom Inventory Short Form, and C) Visual Analog Scale appetite score. In each plot, the top row compares the three timepoints (Start - i.e. Fasted, Mid, and End), and the bottom row plots each of the questions and their responses across all participants. Data shown are mean and standard deviation. Bars are coloured according to kiwifruit variety with maltodextrin shown in blue.

### Blood Vitamin C and Metabolomics

Due to difficulties in blood sampling, only four participants had complete blood datasets (samples collected at fasted and 100 min timepoints across all four visits; eight samples per participant; Figure 5). Three additional participants had partial datasets and were included where possible, while seven participants without control data were excluded. Individual participant data are shown in Figure 5.

**Figure 5:**
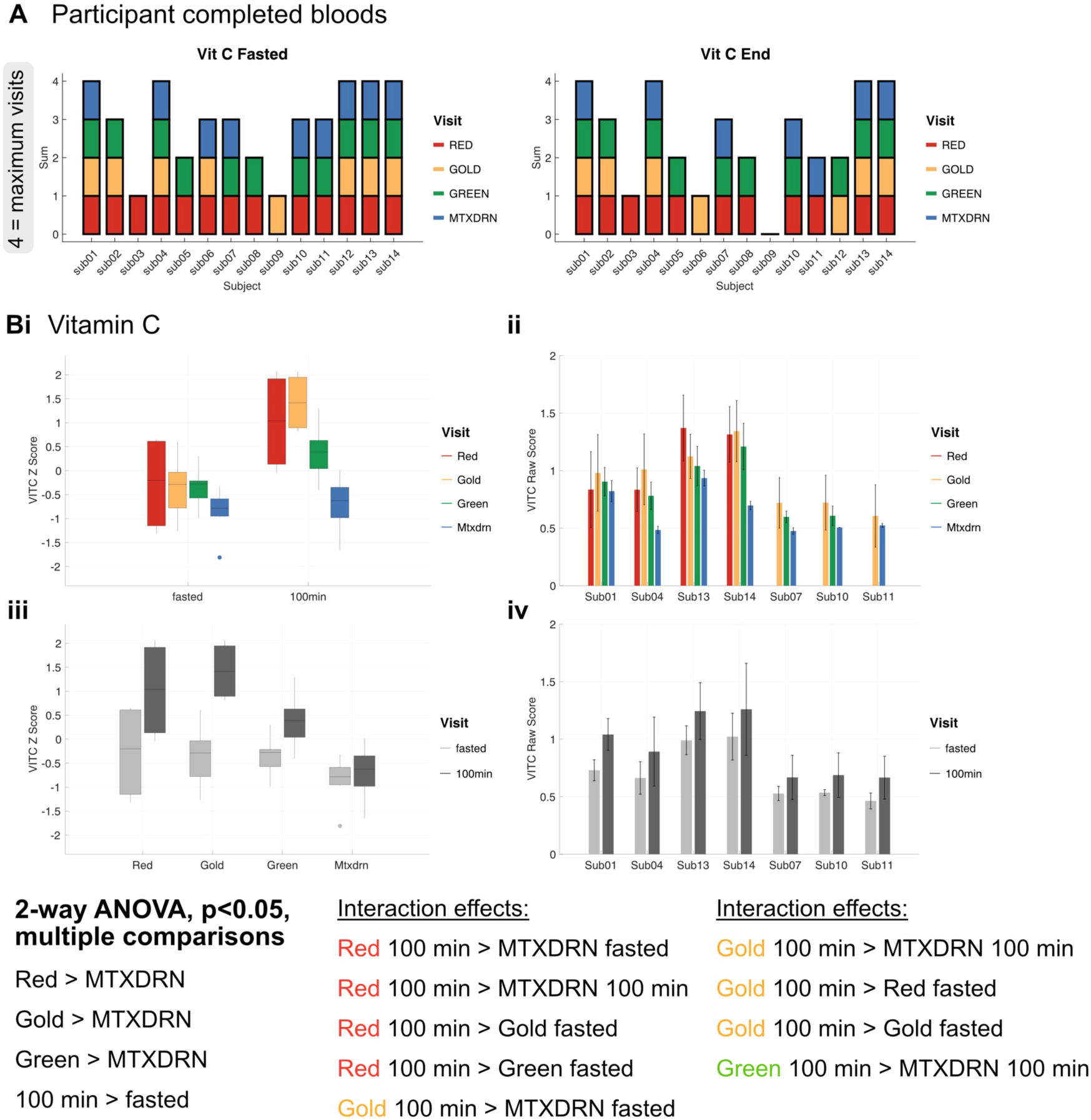
A) Only four participants had complete blood sampling at each visit, 3 other incomplete subjects are also shown. Bi) There were significant differences between kiwifruit and time point for Vitamin C. Bars are coloured according to kiwifruit. Bii shows the results from each participant individually. Biii-iv shows the same data but separated on the x axis by kiwifruit and is coloured according to time point.

Plasma ascorbate (vitamin C) concentrations were within physiologically expected ranges, although inter-individual variability was observed (mean ± SD: fasted 0.85 ± 0.20 mg/100 mL; 100 min 1.11 ± 0.30 mg/100 mL) [23]. A two-way ANOVA (factors: visit [red, green, gold, control] and timepoint [fasted, 100 min]; p < 0.05, corrected) revealed significantly higher ascorbate levels following red, gold, and green kiwifruit compared to the control condition. Ascorbate concentrations were also significantly higher at 100 min compared to the fasted state. A significant interaction effect was observed, with red, gold, and green kiwifruit at 100 min all exceeding control at 100 min.

### N-back fMRI task

#### Behavioural response

Behavioural performance on the N-back task confirmed increasing task difficulty, with the 2-back condition significantly more demanding than the 0-back and 1-back conditions (Figure 6). Reaction time and accuracy measures showed no significant effect of kiwifruit consumption across conditions.

**Figure 6:**
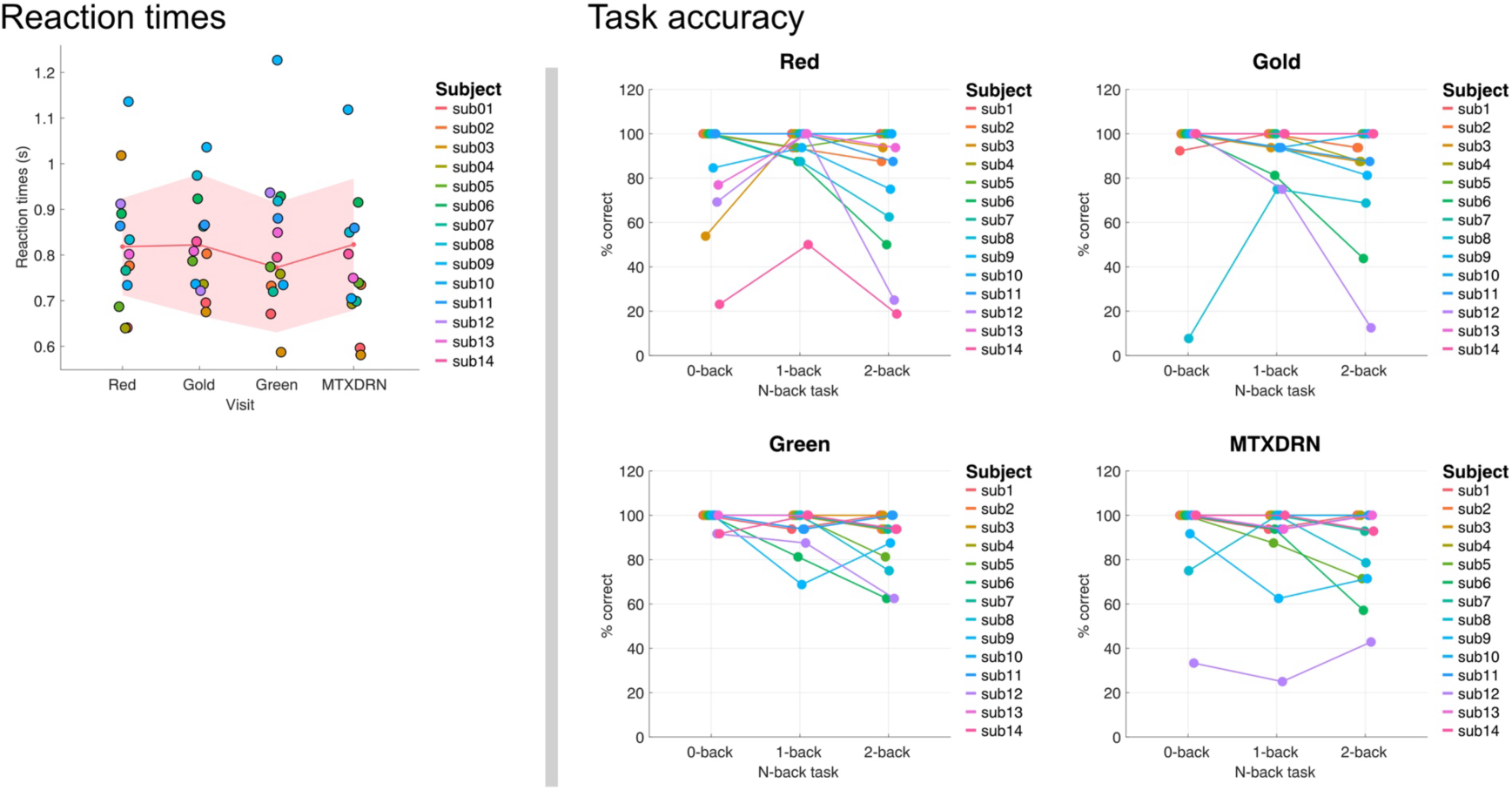
Reaction times and task accuracy across N-back conditions. As expected, performance declined with increasing cognitive load. There were no significant differences between kiwifruit varieties and maltodextrin control.

#### fMRI response

The temporal signal-to-noise ratio (tSNR) of the GE-EPI fMRI data was within the expected range and increased approximately twofold following NORDIC PCA denoising. Robust activation was observed in visual and frontal cortical regions during the N-back task across all conditions (Figure 7). The extent of activation increased with task difficulty. Notably, the maltodextrin condition exhibited more widespread activation compared to the kiwifruit conditions, suggesting reduced neural efficiency. In contrast, red, gold and green kiwifruit were associated with comparatively reduced activation, consistent with lower task demand (Supplementary Table 1).

**Figure 7:**
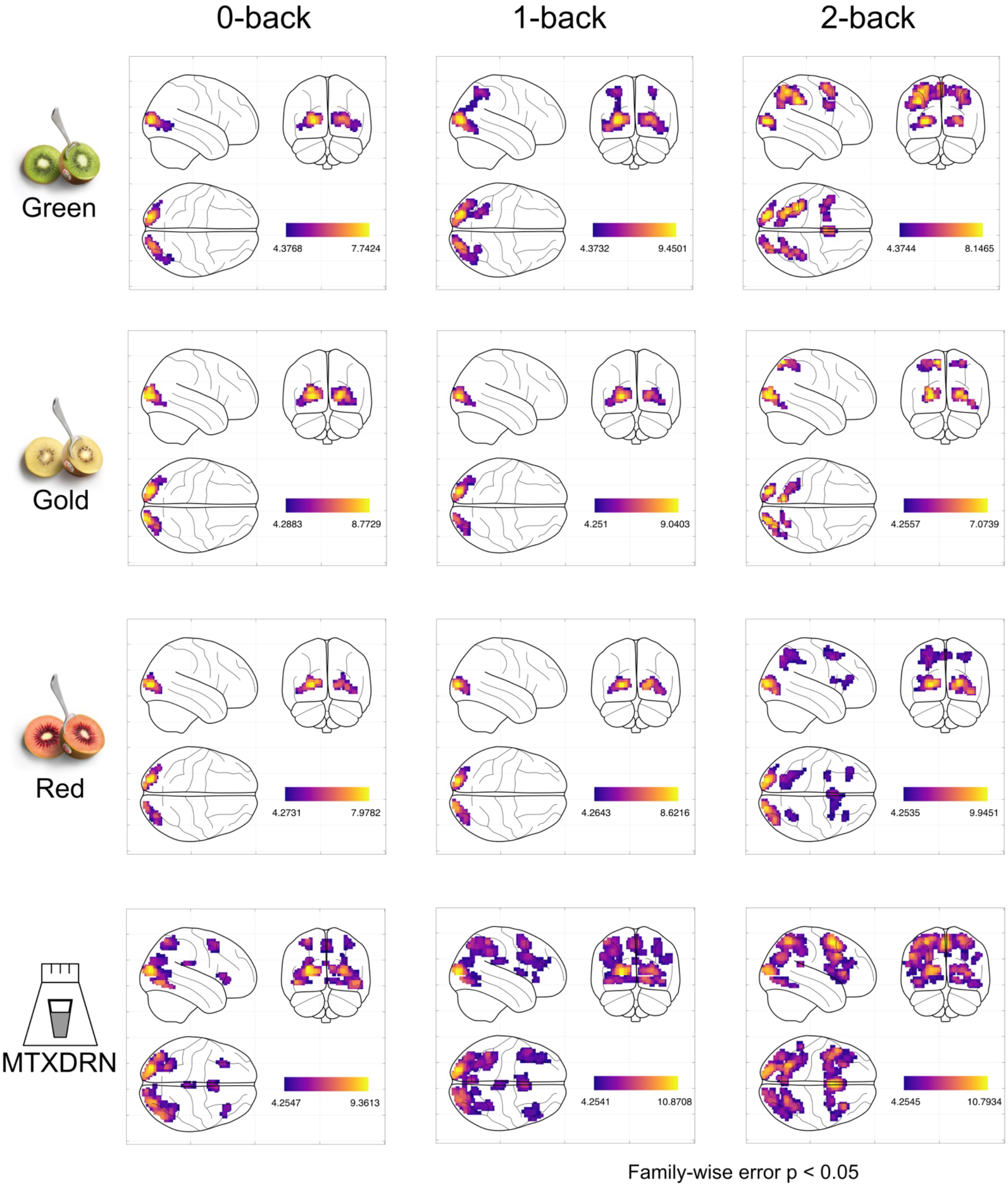
GLM contrast maps for the N-back task following consumption of gold, green, red kiwifruit and maltodextrin control (threshold at p < 0.05, FWE-corrected). Activation extent increased with task difficulty, with the maltodextrin condition showing more global activation than kiwifruit conditions. Note: Less brain areas activated is indicative of less task demand.

The 2-back vs. 0-back contrast was used to isolate brain responses associated with high versus low cognitive load. Significant main effects were observed in regions including the superior frontal gyrus (SFG), anterior and posterior cingulate cortex (ACC, PCC), lateral occipital cortex, and the juxtapositional lobule (supplementary motor area) (Figure 8; Supplementary Table 2).

**Figure 8:**
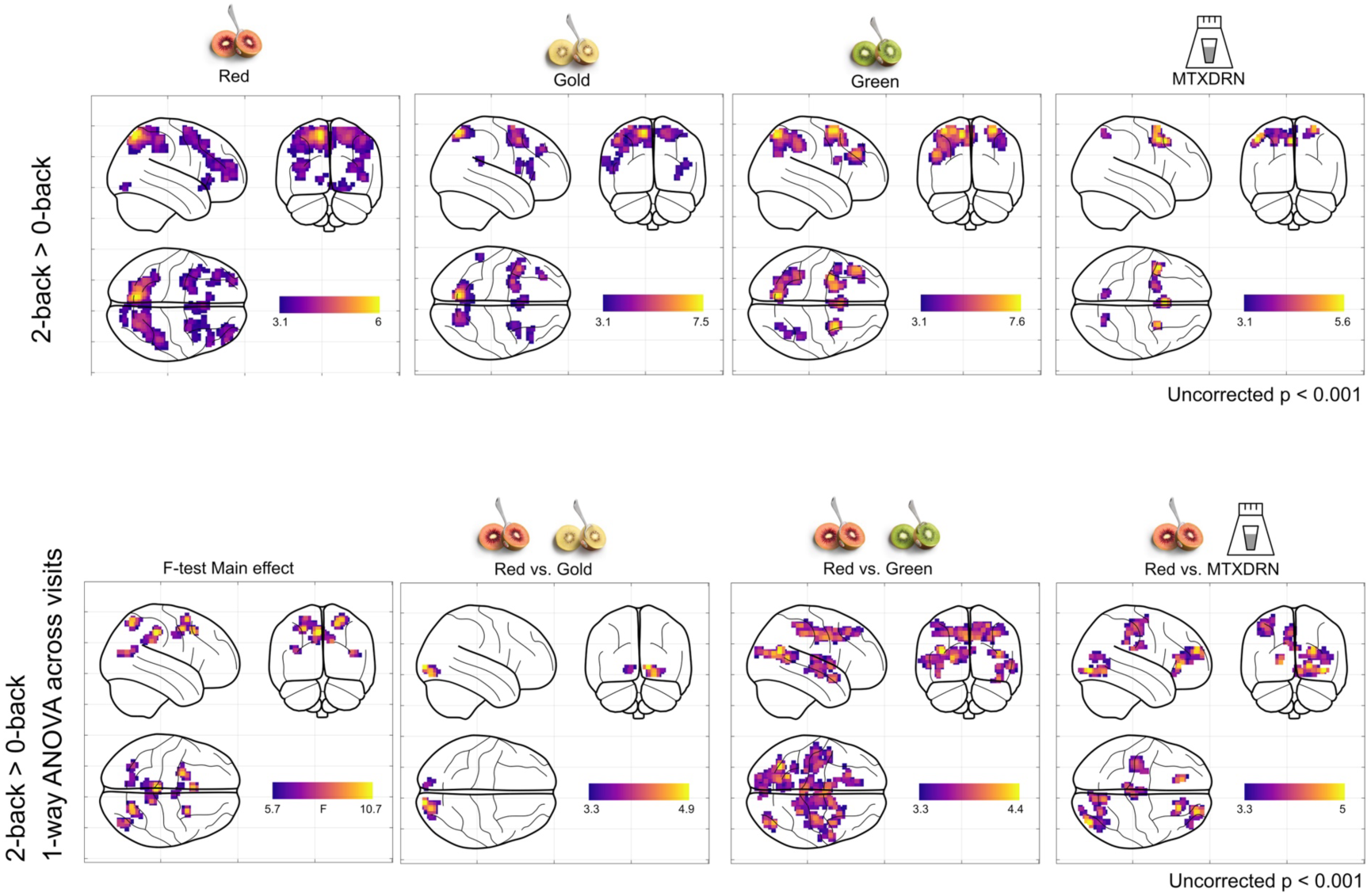
Group-level maps showing the 2-back vs. 0-back contrast following consumption of each kiwifruit across all participants. Bottom row: Second-level analysis comparing the 2-back vs. 0-back contrast between conditions. Significant differences were observed (p < 0.001, uncorrected), with the red kiwifruit condition exhibiting greater activation than the other conditions.

Red kiwifruit elicited greater activation relative to other conditions in key regions involved in working memory and cognitive control, including the ACC, PCC, insula, lateral occipital cortex, pre- and postcentral gyri, and SFG (Supplementary Table 3).

Overall, the maltodextrin condition showed broadly elevated activation across all task conditions, whereas the red kiwifruit condition demonstrated the greatest differentiation between low and high cognitive load, indicating a more targeted and efficient neural response.

### Resting-state fMRI response

Independent component analysis identified differences in salience, sensorimotor (SMN) (red vs. gold), cerebellar, dorsal attention network (DAN) (red vs. green) and DMN networks (red vs. maltodextrin). See Supplementary Figure 1 and the Supplementary Table 4.

## Discussion

This study investigated the acute effects of kiwifruit consumption on cognitive performance, mood, and brain activity using a combination of psychometric, biochemical, and fMRI measures. Overall, the findings suggest that kiwifruit consumption modulates both physiological and neural responses during a working memory task, with effects varying by kiwifruit variety.

### Sensory Evaluation and Implicit Testing Outcomes

Sensory evaluation confirmed that all kiwifruit varieties were well accepted, with no differences in overall liking and no negative emotional associations. As expected, red and gold kiwifruit exhibited higher sugar content (Brix) than green, consistent with their nutritional profiles (green 9.7g/100g, gold 11g/100g and red 11.4g/100g) [24].

PANAS results indicated greater positive affect following kiwifruit consumption com-pared to the maltodextrin control, which may reflect both nutritional effects and differences in sensory experience. The increase in appetite scores over time and the absence of changes in fatigue (MFSI-SF) are consistent with the study duration and the relatively young, healthy participant group.

### Biochemical Findings

Interpretation of blood biomarkers is limited by the small number of complete datasets. Nevertheless, plasma vitamin C levels increased following consumption of all kiwifruit varieties compared to control, consistent with their known vitamin C content (green (88mg/100g), gold (152mg/100g, red (189mg/100g) [25]. The increase in vitamin C between fasted and 100-minute timepoints aligns with established absorption kinetics, where peak plasma concentrations typically occur within 2–3 hours post-ingestion [26].

### N-back fMRI Task

The N-back task elicited expected increases in cognitive load, with the expected reduced performance in task accuracy in the 2-back condition compared to 1-back and 0-back. Brain activation patterns were consistent with established working memory networks, including frontal, parietal, and occipital regions. The lateral occipital cortex is involved in object recognition [27] and was activated primarily in the 0-back condition as the participants focused on a simple task, i.e. recognition of a pre-cued letter, with little working memory component to this condition. Motor regions, such as the juxtapositional lobule (formerly supplementary motor area), were also active; this region has a role in motor planning [28] and in this task, participants were instructed to press a button when they saw the repeated letter.

The 1-back condition did not show an increase in cognitive load for the kiwifruit, but this was clear for the maltodextrin condition, with more frontal regions implicated in memory, spatial task, and error detection, i.e. Superior frontal gyrus, middle frontal gyrus, frontal pole, supramarginal gyrus. For the 2-back task, frontal regions involved in working memory tasks were activated, i.e. dorsolateral prefrontal cortex (frontal cortex, middle frontal gyrus). DLPFC is involved in maintaining and manipulating information in working memory [29]. Regions in the parietal cortex, such as supramarginal gyrus, parietal lobule, are involved in continued involvement of the task. The anterior cingulate cortex (ACC, cingulate gyrus) is involved in error detection, monitoring and cognitive control, and was significantly activated in the 2 vs 0-back contrast [30]. The posterior cingulate gyrus (PCC) was also activated in 2 vs. 0-back; this region is implicated in memory retrieval and self-referential thought [31].

A key observation was the more widespread activation following maltodextrin consumption compared to kiwifruit. This may indicate reduced neural efficiency to the maltodextrin condition; whereby greater brain activation is required to maintain task performance [32]. In contrast, kiwifruit was associated with relatively reduced activation across task conditions. Since maltodextrin is a high-GI carbohydrate, this can lead to a rapid spike in blood sugar levels, followed by a crash, which can negatively impact cognitive performance. Additionally, the poor nutrient composition of maltodextrin compared to kiwifruits may impact efficient brain function. However, there is no consensus in the literature as to the overarching influence of glycaemic index on cognitive functioning [33].

It is important to note that greater activation is not inherently beneficial; what matters is whether neural recruitment is proportional to cognitive demand. In the 2-back vs. 0-back contrast, which isolates high cognitive load, red kiwifruit elicited greater activation in regions implicated in working memory and cognitive control, including the lateral occipital cortex (object recognition), pre and post central gyrus (motor planning, sensory reception), superior frontal gyrus (frontal areas, working memory) and lingual gyrus (visual encoding) [34]. This enhanced differentiation between low and high task demand may reflect a more responsive or efficient neural adaptation to increased cognitive load. The insula was also activated, which has been implicated in working memory tasks, as a processor of emotional information that aids conflict resolution [35]. The above implies that red kiwifruit garnered an enhanced response to difficult portions of the N-back task. This may be due to the higher polyphenol content in red kiwifruit. For the N-back task, the control maltodextrin drink required a greater brain response for lower N-back conditions, possibly implying poorer cognitive efficiency.

### Resting-state fMRI

Group ICA analysis revealed several network regions in the salience and sensorimotor network with greater connectivity for the Red kiwifruit than Gold, Green or maltodextrin. The salience network is important for detecting and filtering relevant stimuli; thus, red kiwifruit may be more attention grabbing than gold kiwifruit [36]. The SMN is involved in processing sensory information and controlling motor actions; greater activation in the precuneus of this network may indicate a stronger self-referential response in the red kiwifruit condition. A region with similar recognised responses related to self-awareness, the SFG of the cerebellar network, was also more active for the Red kiwifruit. There was also increased connectivity in the middle frontal gyrus of the dorsal attention network (DAN). Top-down voluntary processing and focussed attention is a feature of the DAN; the novelty and uniqueness of red kiwifruit may attribute to this [37]. The lateral occipital cortex within the DMN [38] also showed greater connectivity for the Red kiwifruit.

### Potential Mechanisms

While this study did not set out to explore the mechanistic causes of any effects seen following kiwifruit consumption, the observed effects may be driven by the combined action of micronutrients and bioactive compounds present in kiwifruit. Vitamin C plays a role in neurotransmitter synthesis and antioxidant defence, while polyphenols are known to influence cerebral blood flow and neuronal signalling [39,40]. The higher polyphenol and vitamin C content of red kiwifruit may therefore contribute to the greater neural response observed. However, alternative explanations, such as novelty or expectancy effects associated with the red variety, cannot be excluded.

### Limitations and Future Directions

This study has several limitations, including a small sample size and incomplete blood datasets, which limit the generalisability of the biochemical findings. Additionally, the acute design does not capture longer-term effects of regular consumption. The selection of maltodextrin as the control condition was to have an equicaloric comparator to ensure that any observed differences in brain activity could be attributed to the specific properties of the experimental substance, rather than to caloric content. Maltodextrin is a rapidly digestible carbohydrate and may elicit fMRI responses, however, an equicaloric control provides a more physiologically meaningful comparison. Future studies with larger cohorts, controlled blinding, and longitudinal designs are warranted to confirm these findings and further elucidate underlying mechanisms.

## Conclusion

In summary, acute kiwifruit consumption influences mood, metabolic responses, and brain activation during a working memory task, with effects differing by cultivar. Compared to a maltodextrin control, kiwifruit was associated with altered neural responses suggestive of improved cognitive processing. Among the cultivars tested, red kiwifruit demonstrated the most pronounced effects on enhancing cognitive efficiency and modulating brain activity, potentially reflecting its higher micronutrient and polyphenol content.

## CRediT statement

Conceptualization, S.F., S.E., M.A.; methodology, S.F., S.E., M.A. J.H., S.P.; formal analysis, M.A., T.D.; data acquisition, S.F., M.A., S.E.; writing - original draft preparation, S.F., M.A.; writing - review and editing, S.F., M.A., S.E., T.D., J.H.

## Declaration of generative AI use

The authors report generative AI was not used in their research or preparation of this manuscript

## Data Availability Statement

The authors will make this material available upon request to interested researchers.

## Institutional Review Board Statement

The study was conducted in accordance with the Declaration of Helsinki and approved by the Faculty of Medicine and Health Sciences Research Ethics Committee of the University of Nottingham (protocol code FMHS2140223, date of approval 24/04/2023).

## Informed Consent Statement

Informed consent was obtained from all subjects involved in the study.

## Acknowledgements

We thank Rosie Linklater from the Food Experience and Sensory Testing (Feast) Lab, Massey University for her Technical Support during lexicon development, and from the School of Biosciences, University of Nottingham, Yuchen Zheng who helped with the sensory collection and Elizabith Starr who supported kiwi storage.

