## Supplementary Information for "Investigating the effect of acute consumption of kiwifruit on cognitive function using functional MRI"

Participant inclusion and exclusion criteria

Inclusion:

- Aged 18-65
- Participant is willing and able to give informed consent for participation in the study
- Not currently taking any medications that can affect taste perception or appetite (including contraceptive pills)
- Participants with healthy weight (18 Kg/m2 >BMI <25 Kg/m2)
- Ability to understand written and spoken English

Exclusion:

- Allergy to kiwifruit
- Allery to latex
- BMI under 19 or over 25
- Pregnant or breast feeding
- Inability to complete MRI safety questionnaire and / or informed consent process
- Claustrophobia
- Known contraindication to MRI scanning (for example pacemaker / implanted defibrillator, intracranial
- vascular clip, implanted programmable device, intra-ocular metallic fragment, etc)
- Pre-existing taste dysfunction
- Current or history (as declared by the candidate) of major neurological, neurosurgical or psychiatric disorders
- Current or history (as declared by the candidate) of diabetes, heart disease, blood pressure
- Having taken part in a research study in the last 3 months involving invasive procedures or an inconvenience allowance

| Contrast | Network | voxels | p-FWE | p-FDR | p-unc | X | Y | Z | Region |
| --- | --- | --- | --- | --- | --- | --- | --- | --- | --- |
| red_v_gold | Salience | 79 | 0.017 | 0.010 | 0.000 | -10 | 42 | 54 | Frontal Pole |
| red_v_gold | Sensorimotor | 76 | 0.015 | 0.007 | 0.000 | 12 | -64 | 30 | Precuneous Cortex |
| red_v_green | Cerebellar | 53 | 0.085 | 0.049 | 0.001 | 0 | 40 | 50 | Superior Frontal Gyrus |
| red_v_green | Dorsal Attention | 65 | 0.018 | 0.016 | 0.000 | 28 | 34 | 42 | Middle Frontal Gyrus |
| red_v_mtx | Default mode | 96 | 0.001 | 0.001 | 0.000 | 46 | -60 | -4 | Lateral Occipital cortex |

Supplementary Table 4

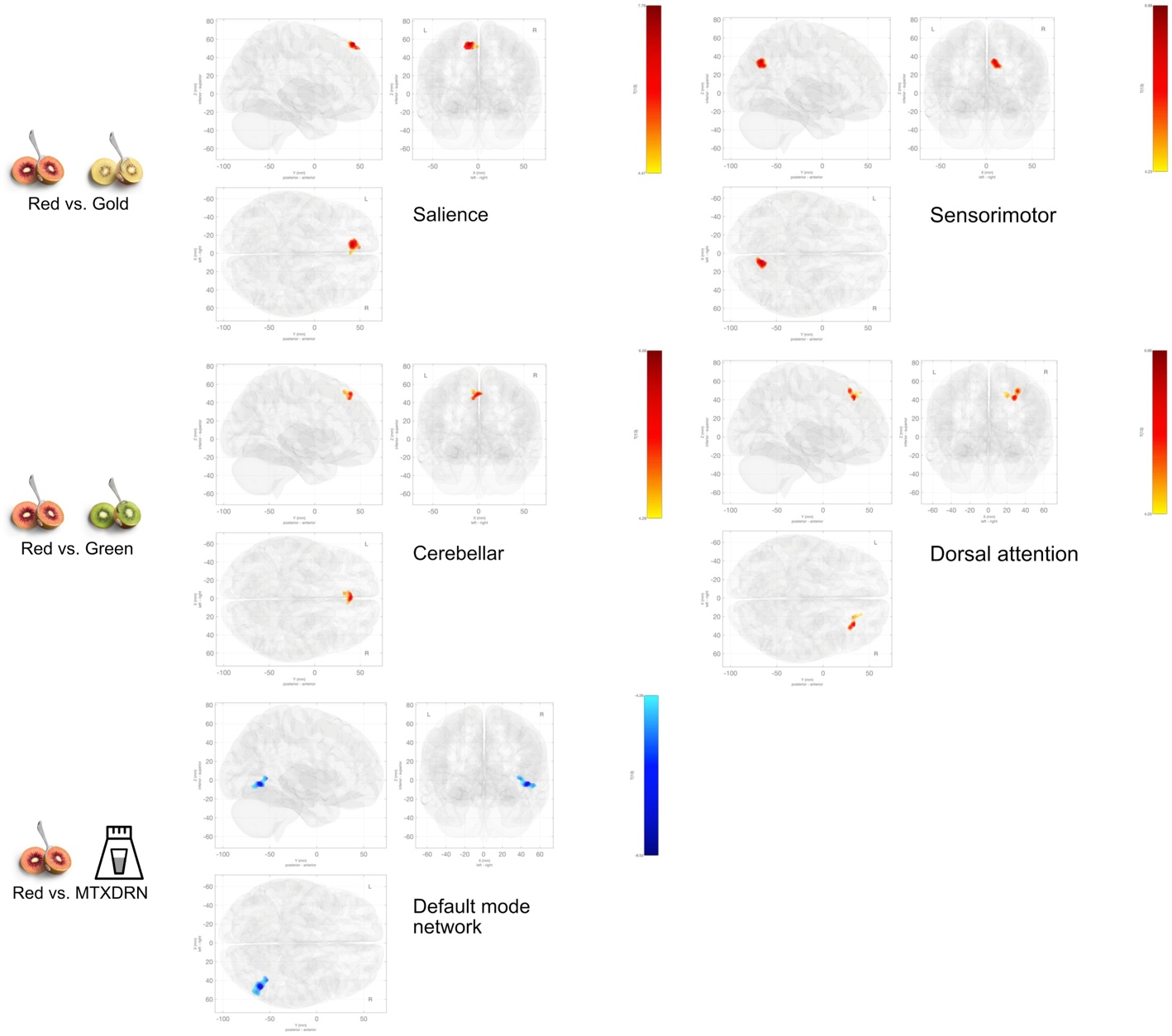

Supplementary Figure 1: Group ICA outputs. Red kiwifruit showed differences between kiwifruit and maltodextrin in several standard networks
