## Supplementary Table 1 for "Investigating the effect of acute consumption of kiwifruit on cognitive function using functional MRI"

| Visit | Contrast | clust_pfwe-corr | clust_qfdr-corr | clust_ke | clust_puncorr | peak_pfwe-corr | peak_qfdr-corr | peak_T | peak_Ze | peak_puncorr | x | y | z | Region |
| --- | --- | --- | --- | --- | --- | --- | --- | --- | --- | --- | --- | --- | --- | --- |
| red | 0-back | 0 | 0.004 | 105 | 0.001 | 0 | 0 | 7.74 | 7.69 | 0 | -24 | -91 | 8 | Occipital Pole |
| red | 0-back |  |  |  |  | 0 | 0 | 6.91 | 6.88 | 0 | -15 | -100 | 11 | Occipital Pole |
| red | 0-back |  |  |  |  | 0.001 | 0.033 | 5.13 | 5.12 | 0 | -36 | -88 | -1 | Lateral Occipital Cortex, inferior division |
| red | 0-back | 0 | 0.01 | 71 | 0.006 | 0 | 0 | 6.24 | 6.22 | 0 | 33 | -82 | 2 | Lateral Occipital Cortex, inferior division |
| red | 0-back |  |  |  |  | 0 | 0 | 6.21 | 6.19 | 0 | 15 | -100 | 5 | Occipital Pole |
| red | 0-back |  |  |  |  | 0 | 0.005 | 5.58 | 5.57 | 0 | 42 | -82 | -4 | Lateral Occipital Cortex, inferior division |
| red | 1-back | 0 | 0 | 175 | 0 | 0 | 0 | 8.89 | inf | 0 | -21 | -94 | 8 | Occipital Pole |
| red | 1-back |  |  |  |  | 0 | 0 | 6.46 | 6.44 | 0 | -36 | -88 | -1 | Lateral Occipital Cortex, inferior division |
| red | 1-back | 0 | 0 | 338 | 0 | 0 | 0 | 8.63 | inf | 0 | 12 | -100 | 2 | Occipital Pole |
| red | 1-back |  |  |  |  | 0 | 0 | 7.63 | 7.59 | 0 | 21 | -97 | 8 | Occipital Pole |
| red | 1-back |  |  |  |  | 0 | 0 | 7.17 | 7.14 | 0 | 21 | -85 | -13 | Occipital Fusiform Gyrus |
| red | 1-back | 0.003 | 0.103 | 30 | 0.059 | 0 | 0 | 6.74 | 6.72 | 0 | -36 | -76 | -13 | Occipital Fusiform Gyrus |
| red | 1-back | 0.002 | 0.103 | 33 | 0.049 | 0 | 0.001 | 5.9 | 5.88 | 0 | -6 | -94 | -7 | Occipital Pole |
| red | 1-back |  |  |  |  | 0 | 0.003 | 5.61 | 5.6 | 0 | -21 | -82 | -13 | Occipital Fusiform Gyrus |
| red | 2-back | 0 | 0 | 255 | 0 | 0 | 0 | 9.83 | inf | 0 | -21 | -94 | 8 | Occipital Pole |
| red | 2-back |  |  |  |  | 0 | 0 | 9.13 | inf | 0 | -15 | -100 | 11 | Occipital Pole |
| red | 2-back |  |  |  |  | 0 | 0 | 6.39 | 6.36 | 0 | -6 | -94 | -7 | Occipital Pole |
| red | 2-back | 0 | 0 | 365 | 0 | 0 | 0 | 9.14 | inf | 0 | 21 | -97 | 11 | Occipital Pole |
| red | 2-back |  |  |  |  | 0 | 0 | 9 | inf | 0 | 12 | -100 | 2 | Occipital Pole |
| red | 2-back |  |  |  |  | 0 | 0 | 8.2 | inf | 0 | 39 | -82 | -4 | Lateral Occipital Cortex, inferior division |
| red | 2-back | 0 | 0.001 | 137 | 0 | 0 | 0 | 6.42 | 6.4 | 0 | -33 | -58 | 53 | Superior Parietal Lobule |
| red | 2-back |  |  |  |  | 0 | 0.006 | 5.62 | 5.6 | 0 | -27 | -70 | 50 | Lateral Occipital Cortex, superior division |
| red | 2-back |  |  |  |  | 0 | 0.013 | 5.43 | 5.42 | 0 | -24 | -64 | 56 | Lateral Occipital Cortex, superior division |
| red | 2-back | 0.001 | 0.058 | 46 | 0.023 | 0.001 | 0.03 | 5.23 | 5.21 | 0 | 0 | 11 | 50 | Paracingulate Gyrus |
| gold | 0-back | 0 | 0 | 478 | 0 | 0 | 0 | 10.37 | inf | 0 | -21 | -94 | 8 | Occipital Pole |
| gold | 0-back |  |  |  |  | 0 | 0 | 7.15 | 7.12 | 0 | -21 | -88 | -10 | Occipital Fusiform Gyrus |
| gold | 0-back |  |  |  |  | 0 | 0 | 7.14 | 7.11 | 0 | -36 | -76 | -13 | Occipital Fusiform Gyrus |
| gold | 0-back | 0 | 0 | 499 | 0 | 0 | 0 | 10.22 | inf | 0 | 21 | -97 | 11 | Occipital Pole |
| gold | 0-back |  |  |  |  | 0 | 0 | 9.5 | inf | 0 | 21 | -85 | -10 | Occipital Fusiform Gyrus |
| gold | 0-back |  |  |  |  | 0 | 0 | 8.62 | inf | 0 | 30 | -85 | 5 | Lateral Occipital Cortex, inferior division |
| gold | 1-back | 0 | 0 | 267 | 0 | 0 | 0 | 9.2 | inf | 0 | -21 | -94 | 8 | Occipital Pole |
| gold | 1-back |  |  |  |  | 0 | 0 | 6.3 | 6.28 | 0 | -42 | -82 | -4 | Lateral Occipital Cortex, inferior division |
| gold | 1-back |  |  |  |  | 0 | 0.001 | 5.97 | 5.95 | 0 | -39 | -73 | -13 | Occipital Fusiform Gyrus |
| gold | 1-back | 0 | 0.001 | 165 | 0 | 0 | 0 | 7.75 | 7.7 | 0 | 15 | -100 | 5 | Occipital Pole |
| gold | 1-back |  |  |  |  | 0 | 0 | 7.64 | 7.6 | 0 | 21 | -97 | 11 | Occipital Pole |
| gold | 1-back |  |  |  |  | 0 | 0 | 6.57 | 6.55 | 0 | 30 | -82 | 5 | Lateral Occipital Cortex, inferior division |
| gold | 1-back | 0.008 | 0.271 | 18 | 0.163 | 0 | 0.001 | 5.77 | 5.76 | 0 | 21 | -85 | -7 | Occipital Fusiform Gyrus |
| gold | 2-back | 0 | 0 | 282 | 0 | 0 | 0 | 8.26 | inf | 0 | -24 | -91 | 14 | Occipital Pole |
| gold | 2-back |  |  |  |  | 0 | 0 | 7.65 | 7.61 | 0 | -39 | -73 | -13 | Occipital Fusiform Gyrus |
| gold | 2-back |  |  |  |  | 0.002 | 0.052 | 5.04 | 5.03 | 0 | -24 | -85 | -10 | Occipital Fusiform Gyrus |
| gold | 2-back | 0 | 0 | 357 | 0 | 0 | 0 | 7.89 | 7.83 | 0 | 24 | -91 | 11 | Occipital Pole |
| gold | 2-back |  |  |  |  | 0 | 0 | 7.6 | 7.56 | 0 | 30 | -85 | 5 | Lateral Occipital Cortex, inferior division |
| gold | 2-back |  |  |  |  | 0 | 0 | 6.64 | 6.61 | 0 | 39 | -85 | -4 | Lateral Occipital Cortex, inferior division |
| gold | 2-back | 0 | 0 | 215 | 0 | 0 | 0 | 7.37 | 7.34 | 0 | -9 | -76 | 53 | Lateral Occipital Cortex, superior division |
| gold | 2-back |  |  |  |  | 0 | 0 | 7.27 | 7.24 | 0 | -12 | -70 | 59 | Lateral Occipital Cortex, superior division |
| gold | 2-back |  |  |  |  | 0 | 0 | 6.48 | 6.46 | 0 | -30 | -61 | 56 | Lateral Occipital Cortex, superior division |
| gold | 2-back | 0.004 | 0.185 | 29 | 0.082 | 0 | 0 | 6.46 | 6.43 | 0 | -51 | 2 | 47 | Precentral Gyrus |
| gold | 2-back |  |  |  |  | 0 | 0.002 | 5.76 | 5.74 | 0 | -36 | -1 | 62 | Middle Frontal Gyrus |
| gold | 2-back |  |  |  |  | 0 | 0.006 | 5.5 | 5.49 | 0 | -42 | 2 | 56 | Middle Frontal Gyrus |
| gold | 2-back | 0 | 0.027 | 82 | 0.007 | 0 | 0 | 6.23 | 6.21 | 0 | 24 | -67 | 59 | Lateral Occipital Cortex, superior division |
| gold | 2-back |  |  |  |  | 0 | 0.001 | 5.87 | 5.86 | 0 | 9 | -73 | 56 | Lateral Occipital Cortex, superior division |
| gold | 2-back |  |  |  |  | 0 | 0.011 | 5.38 | 5.37 | 0 | 39 | -43 | 53 | Superior Parietal Lobule |
| gold | 2-back | 0.005 | 0.185 | 27 | 0.093 | 0 | 0 | 6.1 | 6.08 | 0 | 33 | 26 | 2 | Insular Cortex |
| gold | 2-back | 0.007 | 0.253 | 20 | 0.142 | 0 | 0.002 | 5.81 | 5.8 | 0 | -51 | 14 | 35 | Middle Frontal Gyrus |
| gold | 2-back | 0.001 | 0.077 | 53 | 0.024 | 0 | 0.005 | 5.55 | 5.53 | 0 | -6 | 11 | 47 | Paracingulate Gyrus |
| gold | 2-back |  |  |  |  | 0 | 0.007 | 5.49 | 5.47 | 0 | 0 | 5 | 59 | Juxtapositional Lobule Cortex (formerly Supplementary Motor Cortex) |
| gold | 2-back | 0.009 | 0.278 | 17 | 0.174 | 0.001 | 0.038 | 5.12 | 5.1 | 0 | -27 | -1 | 47 | Middle Frontal Gyrus |
| gold | 2-back | 0.004 | 0.185 | 32 | 0.07 | 0.002 | 0.057 | 5.02 | 5.01 | 0 | -27 | -70 | 32 | Lateral Occipital Cortex, superior division |
| green | 0-back | 0 | 0 | 257 | 0 | 0 | 0 | 8.76 | inf | 0 | -21 | -94 | 8 | Occipital Pole |
| green | 0-back |  |  |  |  | 0 | 0 | 8.09 | inf | 0 | -12 | -100 | 11 | Occipital Pole |
| green | 0-back |  |  |  |  | 0 | 0.002 | 5.89 | 5.87 | 0 | -39 | -76 | -13 | Lateral Occipital Cortex, inferior division |
| green | 0-back | 0 | 0 | 288 | 0 | 0 | 0 | 7.85 | 7.8 | 0 | 21 | -94 | 11 | Occipital Pole |
| green | 0-back |  |  |  |  | 0 | 0 | 7.61 | 7.57 | 0 | 12 | -97 | 8 | Occipital Pole |
| green | 0-back |  |  |  |  | 0 | 0 | 6.93 | 6.89 | 0 | 30 | -85 | 5 | Lateral Occipital Cortex, inferior division |
| green | 1-back | 0 | 0 | 520 | 0 | 0 | 0 | 10.14 | inf | 0 | -27 | -88 | 11 | Lateral Occipital Cortex, superior division |
| green | 1-back |  |  |  |  | 0 | 0 | 9.41 | inf | 0 | -12 | -100 | 11 | Occipital Pole |
| green | 1-back |  |  |  |  | 0 | 0 | 7.64 | 7.59 | 0 | -42 | -79 | -7 | Lateral Occipital Cortex, inferior division |
| green | 1-back | 0 | 0 | 562 | 0 | 0 | 0 | 9.5 | inf | 0 | 21 | -94 | 11 | Occipital Pole |
| green | 1-back |  |  |  |  | 0 | 0 | 9.09 | inf | 0 | 30 | -85 | 5 | Lateral Occipital Cortex, inferior division |
| green | 1-back |  |  |  |  | 0 | 0 | 7.81 | 7.76 | 0 | 21 | -85 | -10 | Occipital Fusiform Gyrus |
| green | 1-back | 0 | 0.005 | 67 | 0.002 | 0 | 0.001 | 6.04 | 6.02 | 0 | -30 | -58 | 53 | Superior Parietal Lobule |
| green | 1-back |  |  |  |  | 0.003 | 0.062 | 5.07 | 5.06 | 0 | -39 | -49 | 53 | Superior Parietal Lobule |
| green | 1-back | 0.001 | 0.047 | 31 | 0.023 | 0 | 0.006 | 5.59 | 5.58 | 0 | -27 | -70 | 29 | Lateral Occipital Cortex, superior division |
| green | 2-back | 0 | 0 | 143 | 0 | 0 | 0 | 8.56 | inf | 0 | -27 | -88 | 8 | Lateral Occipital Cortex, superior division |
| green | 2-back |  |  |  |  | 0 | 0 | 7.58 | 7.54 | 0 | -15 | -97 | 8 | Occipital Pole |
| green | 2-back | 0 | 0 | 462 | 0 | 0 | 0 | 7.96 | inf | 0 | -30 | -58 | 53 | Superior Parietal Lobule |
| green | 2-back |  |  |  |  | 0 | 0 | 7.63 | 7.58 | 0 | -39 | -46 | 50 | Superior Parietal Lobule |
| green | 2-back | 0 | 0 | 153 | 0 | 0 | 0 | 7.57 | 7.53 | 0 | 27 | -85 | 8 | Lateral Occipital Cortex, superior division |
| green | 2-back |  |  |  |  | 0 | 0 | 7.43 | 7.39 | 0 | 12 | -97 | 8 | Occipital Pole |
| green | 2-back | 0 | 0 | 197 | 0 | 0 | 0 | 6.7 | 6.67 | 0 | 27 | -64 | 53 | Lateral Occipital Cortex, superior division |
| green | 2-back |  |  |  |  | 0 | 0 | 6.37 | 6.35 | 0 | 15 | -67 | 50 | Lateral Occipital Cortex, superior division |
| green | 2-back |  |  |  |  | 0 | 0 | 6.2 | 6.18 | 0 | 33 | -46 | 47 | Superior Parietal Lobule |
| green | 2-back | 0 | 0.002 | 80 | 0.001 | 0 | 0 | 6.29 | 6.27 | 0 | 0 | 2 | 62 | Juxtapositional Lobule Cortex (formerly Supplementary Motor Cortex) |
| green | 2-back |  |  |  |  | 0 | 0 | 6.23 | 6.21 | 0 | 0 | 8 | 53 | Paracingulate Gyrus |
| green | 2-back | 0.001 | 0.051 | 30 | 0.025 | 0 | 0.001 | 5.92 | 5.9 | 0 | -36 | -76 | -13 | Occipital Fusiform Gyrus |
| mtxdrn | 0-back | 0 | 0 | 727 | 0 | 0 | 0 | 9.12 | inf | 0 | 18 | -88 | -10 | Occipital Fusiform Gyrus |
| mtxdrn | 0-back |  |  |  |  | 0 | 0 | 8.77 | inf | 0 | 45 | -70 | -13 | Lateral Occipital Cortex, inferior division |
| mtxdrn | 0-back |  |  |  |  | 0 | 0 | 7.96 | inf | 0 | 30 | -85 | 8 | Lateral Occipital Cortex, inferior division |
| mtxdrn | 0-back | 0 | 0 | 597 | 0 | 0 | 0 | 8.3 | inf | 0 | -21 | -91 | 11 | Occipital Pole |
| mtxdrn | 0-back |  |  |  |  | 0 | 0 | 8.01 | inf | 0 | -12 | -100 | 14 | Occipital Pole |
| mtxdrn | 0-back |  |  |  |  | 0 | 0 | 7.88 | 7.83 | 0 | -42 | -76 | -10 | Lateral Occipital Cortex, inferior division |
| mtxdrn | 0-back | 0.001 | 0.104 | 62 | 0.026 | 0 | 0 | 6.05 | 6.03 | 0 | -30 | -61 | 56 | Lateral Occipital Cortex, superior division |
| mtxdrn | 0-back |  |  |  |  | 0.004 | 0.105 | 4.85 | 4.84 | 0 | -27 | -67 | 47 | Lateral Occipital Cortex, superior division |
| mtxdrn | 0-back | 0.002 | 0.138 | 48 | 0.046 | 0 | 0.001 | 5.93 | 5.91 | 0 | 3 | 8 | 53 | Juxtapositional Lobule Cortex (formerly Supplementary Motor Cortex) |
| mtxdrn | 0-back | 0.01 | 0.381 | 19 | 0.19 | 0 | 0.004 | 5.59 | 5.58 | 0 | -33 | 20 | 5 | Insular Cortex |
| mtxdrn | 0-back | 0.006 | 0.266 | 29 | 0.111 | 0.002 | 0.053 | 5.04 | 5.02 | 0 | 33 | -55 | 56 | Superior Parietal Lobule |
| mtxdrn | 0-back |  |  |  |  | 0.031 | 0.632 | 4.37 | 4.36 | 0 | 30 | -58 | 47 | Lateral Occipital Cortex, superior division |
| mtxdrn | 1-back | 0 | 0 | 1604 | 0 | 0 | 0 | 11.77 | inf | 0 | 21 | -82 | -13 | Occipital Fusiform Gyrus |
| mtxdrn | 1-back |  |  |  |  | 0 | 0 | 11.33 | inf | 0 | -24 | -82 | -13 | Occipital Fusiform Gyrus |
| mtxdrn | 1-back |  |  |  |  | 0 | 0 | 10.94 | inf | 0 | -21 | -94 | 11 | Occipital Pole |
| mtxdrn | 1-back | 0 | 0.002 | 215 | 0 | 0 | 0 | 8.21 | inf | 0 | 3 | 8 | 53 | Juxtapositional Lobule Cortex (formerly Supplementary Motor Cortex) |
| mtxdrn | 1-back |  |  |  |  | 0 | 0 | 7.01 | 6.98 | 0 | 3 | 2 | 65 | Juxtapositional Lobule Cortex (formerly Supplementary Motor Cortex) |
| mtxdrn | 1-back | 0 | 0 | 311 | 0 | 0 | 0 | 7.71 | 7.67 | 0 | -45 | -49 | 56 | Supramarginal Gyrus, posterior division |
| mtxdrn | 1-back |  |  |  |  | 0 | 0 | 6.53 | 6.51 | 0 | -36 | -46 | 41 | Supramarginal Gyrus, posterior division |
| mtxdrn | 1-back |  |  |  |  | 0 | 0 | 6.46 | 6.44 | 0 | -27 | -52 | 44 | Superior Parietal Lobule |
| mtxdrn | 1-back | 0 | 0.02 | 117 | 0.004 | 0 | 0 | 7.52 | 7.48 | 0 | -6 | -76 | 50 | Precuneous Cortex |
| mtxdrn | 1-back |  |  |  |  | 0 | 0.014 | 5.35 | 5.34 | 0 | 12 | -70 | 47 | Precuneous Cortex |
| mtxdrn | 1-back |  |  |  |  | 0 | 0.016 | 5.33 | 5.32 | 0 | 12 | -67 | 38 | Precuneous Cortex |
| mtxdrn | 1-back | 0.001 | 0.081 | 71 | 0.018 | 0 | 0 | 6.65 | 6.63 | 0 | 0 | -28 | 26 | Cingulate Gyrus, posterior division |
| mtxdrn | 1-back |  |  |  |  | 0.006 | 0.152 | 4.77 | 4.76 | 0 | 6 | -16 | 26 | Cingulate Gyrus, posterior division |
| mtxdrn | 1-back | 0 | 0.015 | 131 | 0.002 | 0 | 0 | 6.42 | 6.4 | 0 | 33 | -64 | 56 | Lateral Occipital Cortex, superior division |
| mtxdrn | 1-back |  |  |  |  | 0 | 0.005 | 5.64 | 5.63 | 0 | 30 | -46 | 44 | Superior Parietal Lobule |
| mtxdrn | 1-back |  |  |  |  | 0 | 0.006 | 5.55 | 5.54 | 0 | 36 | -52 | 53 | Superior Parietal Lobule |
| mtxdrn | 1-back | 0.009 | 0.348 | 20 | 0.18 | 0 | 0.001 | 5.89 | 5.87 | 0 | -9 | 23 | 29 | Cingulate Gyrus, anterior division |
| mtxdrn | 1-back | 0.004 | 0.269 | 35 | 0.083 | 0 | 0.004 | 5.66 | 5.64 | 0 | -42 | -4 | 59 | Precentral Gyrus |
| mtxdrn | 1-back |  |  |  |  | 0.002 | 0.057 | 5.01 | 5 | 0 | -27 | -4 | 50 | Middle Frontal Gyrus |
| mtxdrn | 1-back | 0.007 | 0.314 | 24 | 0.144 | 0 | 0.005 | 5.63 | 5.61 | 0 | 27 | 62 | -1 | Frontal Pole |
| mtxdrn | 1-back |  |  |  |  | 0.003 | 0.072 | 4.96 | 4.94 | 0 | 21 | 50 | 2 | Frontal Pole |
| mtxdrn | 1-back | 0 | 0.013 | 144 | 0.002 | 0 | 0.005 | 5.6 | 5.59 | 0 | -48 | 14 | 32 | Middle Frontal Gyrus |
| mtxdrn | 1-back |  |  |  |  | 0 | 0.014 | 5.35 | 5.34 | 0 | -36 | 32 | 17 | Inferior Frontal Gyrus, pars triangularis |
| mtxdrn | 1-back |  |  |  |  | 0.001 | 0.036 | 5.13 | 5.11 | 0 | -42 | 2 | 35 | Precentral Gyrus |
| mtxdrn | 1-back | 0.002 | 0.151 | 52 | 0.039 | 0 | 0.013 | 5.4 | 5.38 | 0 | 54 | 20 | 32 | Middle Frontal Gyrus |
| mtxdrn | 1-back |  |  |  |  | 0.002 | 0.05 | 5.04 | 5.03 | 0 | 45 | 14 | 29 | Inferior Frontal Gyrus, pars opercularis |
| mtxdrn | 1-back |  |  |  |  | 0.003 | 0.093 | 4.89 | 4.88 | 0 | 45 | 23 | 29 | Middle Frontal Gyrus |
| mtxdrn | 1-back | 0.005 | 0.269 | 32 | 0.096 | 0 | 0.013 | 5.39 | 5.38 | 0 | -33 | 20 | 8 | Frontal Opercular Cortex |
| mtxdrn | 1-back | 0.006 | 0.314 | 27 | 0.123 | 0 | 0.017 | 5.31 | 5.3 | 0 | 36 | 23 | 8 | Frontal Opercular Cortex |
| mtxdrn | 1-back |  |  |  |  | 0.021 | 0.474 | 4.47 | 4.46 | 0 | 36 | 26 | -4 | Frontal Orbital Cortex |
| mtxdrn | 1-back | 0.007 | 0.314 | 24 | 0.144 | 0.001 | 0.033 | 5.15 | 5.14 | 0 | 45 | 2 | 53 | Middle Frontal Gyrus |
| mtxdrn | 2-back | 0 | 0 | 240 | 0 | 0 | 0 | 10.49 | inf | 0 | 0 | 5 | 56 | Juxtapositional Lobule Cortex (formerly Supplementary Motor Cortex) |
| mtxdrn | 2-back |  |  |  |  | 0 | 0 | 10.15 | inf | 0 | 0 | 11 | 50 | Paracingulate Gyrus |
| mtxdrn | 2-back | 0 | 0 | 460 | 0 | 0 | 0 | 9.86 | inf | 0 | 18 | -85 | -10 | Occipital Fusiform Gyrus |
| mtxdrn | 2-back |  |  |  |  | 0 | 0 | 9.83 | inf | 0 | 15 | -94 | -1 | Occipital Pole |
| mtxdrn | 2-back |  |  |  |  | 0 | 0 | 7.89 | 7.84 | 0 | 24 | -91 | 11 | Occipital Pole |
| mtxdrn | 2-back | 0 | 0 | 492 | 0 | 0 | 0 | 9.24 | inf | 0 | -21 | -94 | 8 | Occipital Pole |
| mtxdrn | 2-back |  |  |  |  | 0 | 0 | 7.8 | 7.75 | 0 | -42 | -73 | -7 | Lateral Occipital Cortex, inferior division |
| mtxdrn | 2-back |  |  |  |  | 0 | 0 | 7.79 | 7.74 | 0 | -6 | -94 | -4 | Occipital Pole |
| mtxdrn | 2-back | 0 | 0 | 359 | 0 | 0 | 0 | 8.76 | inf | 0 | -30 | -61 | 56 | Lateral Occipital Cortex, superior division |
| mtxdrn | 2-back |  |  |  |  | 0 | 0 | 7.18 | 7.15 | 0 | -24 | -64 | 62 | Lateral Occipital Cortex, superior division |
| mtxdrn | 2-back |  |  |  |  | 0 | 0 | 7.17 | 7.14 | 0 | -27 | -70 | 29 | Lateral Occipital Cortex, superior division |
| mtxdrn | 2-back | 0 | 0 | 243 | 0 | 0 | 0 | 8.34 | inf | 0 | -54 | 11 | 35 | Middle Frontal Gyrus |
| mtxdrn | 2-back |  |  |  |  | 0 | 0 | 8.22 | inf | 0 | -48 | 5 | 50 | Middle Frontal Gyrus |
| mtxdrn | 2-back |  |  |  |  | 0 | 0 | 8 | inf | 0 | -42 | 2 | 56 | Middle Frontal Gyrus |
| mtxdrn | 2-back | 0 | 0.001 | 206 | 0 | 0 | 0 | 8.3 | inf | 0 | 27 | -64 | 56 | Lateral Occipital Cortex, superior division |
| mtxdrn | 2-back |  |  |  |  | 0 | 0 | 6.85 | 6.82 | 0 | 33 | -49 | 56 | Superior Parietal Lobule |
| mtxdrn | 2-back |  |  |  |  | 0.001 | 0.029 | 5.14 | 5.12 | 0 | 30 | -46 | 47 | Superior Parietal Lobule |
| mtxdrn | 2-back | 0.003 | 0.135 | 45 | 0.052 | 0 | 0 | 6.74 | 6.71 | 0 | -30 | 23 | 5 | Insular Cortex |
| mtxdrn | 2-back | 0.009 | 0.339 | 21 | 0.17 | 0 | 0.001 | 5.81 | 5.79 | 0 | 33 | 26 | 5 | Frontal Opercular Cortex |
| mtxdrn | 2-back | 0.007 | 0.291 | 26 | 0.129 | 0 | 0.004 | 5.57 | 5.56 | 0 | 24 | 2 | 62 | Superior Frontal Gyrus |
| mtxdrn | 2-back |  |  |  |  | 0.042 | 0.833 | 4.29 | 4.28 | 0 | 30 | -1 | 56 | Middle Frontal Gyrus |
| mtxdrn | 2-back | 0.012 | 0.414 | 15 | 0.242 | 0 | 0.008 | 5.41 | 5.4 | 0 | 51 | 5 | 35 | Precentral Gyrus |
