## Supplementary Table 2 for "Investigating the effect of acute consumption of kiwifruit on cognitive function using functional MRI"

| Visit | clust_pfwe-corr | clust_qfdr-corr | clust_ke | clust_puncorr | peak_pfwe-corr | peak_qfdr-corr | peak_T | peak_Ze | peak_puncorr | x | y | z | Region |
| --- | --- | --- | --- | --- | --- | --- | --- | --- | --- | --- | --- | --- | --- |
| red | 0 | 0.002 | 489 | 0 | 0 | 0 | 7 | 6.97 | 0 | -12 | -70 | 59 | Lateral Occipital Cortex, superior division |
| red |  |  |  |  | 0.002 | 0.018 | 5.12 | 5.11 | 0 | -33 | -58 | 53 | Superior Parietal Lobule |
| red |  |  |  |  | 0.109 | 0.165 | 4.09 | 4.08 | 0 | -36 | -43 | 41 | Superior Parietal Lobule |
| red | 0 | 0.002 | 467 | 0 | 0.002 | 0.018 | 5.11 | 5.1 | 0 | 24 | -67 | 56 | Lateral Occipital Cortex, superior division |
| red |  |  |  |  | 0.008 | 0.053 | 4.76 | 4.76 | 0 | 12 | -61 | 65 | Lateral Occipital Cortex, superior division |
| red |  |  |  |  | 0.008 | 0.053 | 4.76 | 4.75 | 0 | 45 | -40 | 53 | Supramarginal Gyrus, posterior division |
| red | 0.063 | 0.214 | 119 | 0.027 | 0.017 | 0.08 | 4.58 | 4.58 | 0 | 45 | 47 | 2 | Frontal Pole |
| red |  |  |  |  | 0.041 | 0.127 | 4.36 | 4.35 | 0 | 39 | 50 | 20 | Frontal Pole |
| red |  |  |  |  | 0.085 | 0.151 | 4.16 | 4.15 | 0 | 30 | 47 | 11 | Frontal Pole |
| red | 0.027 | 0.15 | 164 | 0.011 | 0.021 | 0.088 | 4.53 | 4.52 | 0 | -30 | -4 | 65 | Superior Frontal Gyrus |
| red |  |  |  |  | 0.064 | 0.133 | 4.24 | 4.23 | 0 | -42 | 8 | 50 | Middle Frontal Gyrus |
| red |  |  |  |  | 0.094 | 0.151 | 4.13 | 4.13 | 0 | -21 | 2 | 59 | Superior Frontal Gyrus |
| red | 0.057 | 0.214 | 124 | 0.024 | 0.033 | 0.121 | 4.42 | 4.41 | 0 | -33 | 56 | 17 | Frontal Pole |
| red |  |  |  |  | 0.059 | 0.133 | 4.26 | 4.25 | 0 | -39 | 35 | 17 | Middle Frontal Gyrus |
| red |  |  |  |  | 0.066 | 0.133 | 4.23 | 4.22 | 0 | -36 | 56 | 5 | Frontal Pole |
| red | 0.147 | 0.371 | 78 | 0.065 | 0.042 | 0.127 | 4.35 | 4.34 | 0 | 0 | 17 | 41 | Paracingulate Gyrus |
| red |  |  |  |  | 0.331 | 0.338 | 3.73 | 3.72 | 0 | 0 | 5 | 56 | Juxtapositional Lobule Cortex (formerly Supplementary Motor Cortex) |
| red | 0.121 | 0.352 | 87 | 0.053 | 0.049 | 0.127 | 4.31 | 4.31 | 0 | 27 | -1 | 65 | Superior Frontal Gyrus |
| red |  |  |  |  | 0.159 | 0.189 | 3.98 | 3.97 | 0 | 21 | 2 | 53 | Superior Frontal Gyrus |
| red |  |  |  |  | 0.575 | 0.511 | 3.48 | 3.47 | 0 | 42 | 8 | 50 | Middle Frontal Gyrus |
| red | 0.481 | 0.861 | 26 | 0.269 | 0.089 | 0.151 | 4.15 | 4.14 | 0 | 51 | 26 | 32 | Middle Frontal Gyrus |
| red |  |  |  |  | 0.173 | 0.195 | 3.95 | 3.94 | 0 | 42 | 26 | 29 | Middle Frontal Gyrus |
| red | 0.516 | 0.861 | 23 | 0.298 | 0.143 | 0.188 | 4.01 | 4 | 0 | 18 | -85 | -13 | Occipital Fusiform Gyrus |
| red |  |  |  |  | 0.388 | 0.378 | 3.66 | 3.66 | 0 | 3 | -82 | -7 | Lingual Gyrus |
| red | 0.582 | 0.861 | 18 | 0.358 | 0.566 | 0.51 | 3.49 | 3.48 | 0 | 27 | 20 | -10 | Frontal Orbital Cortex |
| gold | 0.04 | 0.294 | 156 | 0.019 | 0 | 0 | 8.33 | inf | 0 | -12 | -70 | 59 | Lateral Occipital Cortex, superior division |
| gold |  |  |  |  | 0.447 | 0.405 | 3.56 | 3.56 | 0 | -9 | -55 | 56 | Precuneous Cortex |
| gold |  |  |  |  | 0.53 | 0.483 | 3.48 | 3.48 | 0 | 6 | -61 | 59 | Precuneous Cortex |
| gold | 0.046 | 0.294 | 148 | 0.022 | 0 | 0 | 6.41 | 6.39 | 0 | -45 | 8 | 50 | Middle Frontal Gyrus |
| gold |  |  |  |  | 0 | 0 | 5.88 | 5.86 | 0 | -30 | -1 | 65 | Middle Frontal Gyrus |
| gold |  |  |  |  | 0 | 0.002 | 5.38 | 5.37 | 0 | -42 | 2 | 56 | Middle Frontal Gyrus |
| gold | 0.151 | 0.681 | 82 | 0.076 | 0 | 0.002 | 5.4 | 5.38 | 0 | 15 | -70 | 59 | Lateral Occipital Cortex, superior division |
| gold | 0.572 | 0.872 | 18 | 0.391 | 0.001 | 0.003 | 5.17 | 5.16 | 0 | -36 | 38 | 38 | Middle Frontal Gyrus |
| gold | 0.254 | 0.872 | 56 | 0.135 | 0.027 | 0.043 | 4.43 | 4.43 | 0 | -6 | 14 | 44 | Paracingulate Gyrus |
| gold |  |  |  |  | 0.076 | 0.097 | 4.16 | 4.16 | 0 | 0 | 8 | 62 | Juxtapositional Lobule Cortex (formerly Supplementary Motor Cortex) |
| gold | 0.43 | 0.872 | 31 | 0.259 | 0.031 | 0.044 | 4.4 | 4.4 | 0 | -51 | 5 | 11 | Precentral Gyrus |
| gold |  |  |  |  | 0.225 | 0.23 | 3.83 | 3.82 | 0 | -45 | 5 | 20 | Precentral Gyrus |
| gold |  |  |  |  | 0.28 | 0.271 | 3.75 | 3.75 | 0 | -57 | 11 | 17 | Inferior Frontal Gyrus, pars opercularis |
| gold | 0.3 | 0.872 | 48 | 0.164 | 0.049 | 0.067 | 4.28 | 4.27 | 0 | 27 | 2 | 59 | Superior Frontal Gyrus |
| gold |  |  |  |  | 0.195 | 0.205 | 3.88 | 3.87 | 0 | 36 | 8 | 59 | Middle Frontal Gyrus |
| gold | 0.412 | 0.872 | 33 | 0.245 | 0.092 | 0.111 | 4.11 | 4.1 | 0 | 42 | 23 | 17 | Inferior Frontal Gyrus, pars triangularis |
| gold |  |  |  |  | 0.122 | 0.135 | 4.02 | 4.02 | 0 | 33 | 26 | 2 | Insular Cortex |
| gold |  |  |  |  | 0.328 | 0.315 | 3.69 | 3.69 | 0 | 33 | 20 | 11 | Frontal Opercular Cortex |
| gold | 0.572 | 0.872 | 18 | 0.391 | 0.096 | 0.111 | 4.09 | 4.09 | 0 | -63 | -43 | 23 | Supramarginal Gyrus, posterior division |
| green | 0 | 0.001 | 405 | 0 | 0 | 0 | 6.27 | 6.25 | 0 | -12 | -70 | 56 | Lateral Occipital Cortex, superior division |
| green |  |  |  |  | 0.004 | 0.005 | 4.98 | 4.97 | 0 | -24 | -64 | 59 | Lateral Occipital Cortex, superior division |
| green |  |  |  |  | 0.004 | 0.005 | 4.98 | 4.96 | 0 | -36 | -43 | 44 | Superior Parietal Lobule |
| green | 0.078 | 0.074 | 88 | 0.025 | 0 | 0 | 6.13 | 6.11 | 0 | 30 | 5 | 62 | Superior Frontal Gyrus |
| green | 0.022 | 0.05 | 138 | 0.007 | 0 | 0 | 6.09 | 6.07 | 0 | -33 | -1 | 62 | Middle Frontal Gyrus |
| green |  |  |  |  | 0 | 0 | 5.61 | 5.59 | 0 | -21 | 5 | 62 | Superior Frontal Gyrus |
| green |  |  |  |  | 0.812 | 0.612 | 3.35 | 3.35 | 0 | -42 | 2 | 56 | Middle Frontal Gyrus |
| green | 0.058 | 0.074 | 99 | 0.018 | 0 | 0 | 5.58 | 5.57 | 0 | -45 | 35 | 29 | Middle Frontal Gyrus |
| green |  |  |  |  | 0.131 | 0.08 | 4.12 | 4.11 | 0 | -33 | 38 | 38 | Middle Frontal Gyrus |
| green | 0.078 | 0.074 | 88 | 0.025 | 0.001 | 0.002 | 5.29 | 5.27 | 0 | 0 | 11 | 56 | Superior Frontal Gyrus |
| green |  |  |  |  | 0.002 | 0.003 | 5.12 | 5.11 | 0 | -3 | 14 | 47 | Paracingulate Gyrus |
| green |  |  |  |  | 0.047 | 0.036 | 4.4 | 4.39 | 0 | -3 | 2 | 65 | Juxtapositional Lobule Cortex (formerly Supplementary Motor Cortex) |
| green | 0.532 | 0.431 | 22 | 0.23 | 0.059 | 0.039 | 4.34 | 4.33 | 0 | -45 | 8 | 23 | Inferior Frontal Gyrus, pars opercularis |
| green | 0.339 | 0.268 | 37 | 0.125 | 0.105 | 0.067 | 4.18 | 4.18 | 0 | 42 | -40 | 47 | Supramarginal Gyrus, posterior division |
| green |  |  |  |  | 0.761 | 0.571 | 3.41 | 3.4 | 0 | 33 | -43 | 44 | Superior Parietal Lobule |
| green | 0.319 | 0.268 | 39 | 0.116 | 0.172 | 0.102 | 4.04 | 4.03 | 0 | 39 | -55 | 53 | Angular Gyrus |
| green |  |  |  |  | 0.333 | 0.19 | 3.82 | 3.81 | 0 | 30 | -64 | 50 | Lateral Occipital Cortex, superior division |
| green |  |  |  |  | 0.627 | 0.44 | 3.53 | 3.53 | 0 | 27 | -70 | 56 | Lateral Occipital Cortex, superior division |
| mtxdrn | 0.173 | 0.734 | 81 | 0.101 | 0 | 0.001 | 5.42 | 5.41 | 0 | -48 | 5 | 50 | Middle Frontal Gyrus |
| mtxdrn |  |  |  |  | 0.041 | 0.063 | 4.29 | 4.28 | 0 | -24 | -1 | 47 | Superior Frontal Gyrus |
| mtxdrn |  |  |  |  | 0.14 | 0.15 | 3.94 | 3.94 | 0 | -21 | 5 | 59 | Superior Frontal Gyrus |
| mtxdrn | 0.265 | 0.734 | 57 | 0.162 | 0 | 0.001 | 5.39 | 5.38 | 0 | -3 | 14 | 47 | Paracingulate Gyrus |
| mtxdrn |  |  |  |  | 0.241 | 0.24 | 3.76 | 3.76 | 0 | 0 | 5 | 62 | Juxtapositional Lobule Cortex (formerly Supplementary Motor Cortex) |
| mtxdrn | 0.546 | 0.734 | 19 | 0.417 | 0.002 | 0.004 | 5.07 | 5.05 | 0 | 27 | 5 | 65 | Superior Frontal Gyrus |
| mtxdrn | 0.504 | 0.734 | 23 | 0.371 | 0.116 | 0.15 | 4 | 3.99 | 0 | 21 | -67 | 62 | Lateral Occipital Cortex, superior division |
| mtxdrn | 0.557 | 0.734 | 18 | 0.431 | 0.137 | 0.15 | 3.95 | 3.94 | 0 | -12 | -64 | 56 | Lateral Occipital Cortex, superior division |
