## Supplementary Table 3 for "Investigating the effect of acute consumption of kiwifruit on cognitive function using functional MRI"

| Contrast | clust_pfwe-corr | clust_qfdr-corr | clust_ke | clust_puncorr | peak_pfwe-corr | peak_qfdr-corr | peak_T | peak_Ze | peak_puncorr | x | y | z | Region |
| --- | --- | --- | --- | --- | --- | --- | --- | --- | --- | --- | --- | --- | --- |
| red_v_green | 0 | 0 | 272 | 0 | 0.083 | 0.519 | 5.11 | 4.44 | 0 | -36 | -64 | 20 | Lateral Occipital Cortex, superior division |
| red_v_green |  |  |  |  | 0.146 | 0.519 | 4.88 | 4.28 | 0 | -21 | -55 | 11 | Precuneous Cortex |
| red_v_green | 0.066 | 0.125 | 55 | 0.011 | 0.171 | 0.519 | 4.81 | 4.24 | 0 | 39 | -73 | 17 | Lateral Occipital Cortex, superior division |
| red_v_green |  |  |  |  | 0.278 | 0.519 | 4.59 | 4.08 | 0 | 39 | -82 | 17 | Lateral Occipital Cortex, superior division |
| red_v_green | 0 | 0 | 451 | 0 | 0.217 | 0.519 | 4.71 | 4.16 | 0 | -3 | -4 | 38 | Cingulate Gyrus, anterior division |
| red_v_green |  |  |  |  | 0.259 | 0.519 | 4.63 | 4.11 | 0 | 33 | -19 | 44 | Precentral Gyrus |
| red_v_green | 0.46 | 0.546 | 20 | 0.102 | 0.302 | 0.519 | 4.55 | 4.05 | 0 | 39 | 5 | -16 | Insular Cortex |
| red_v_green | 0.197 | 0.303 | 35 | 0.036 | 0.337 | 0.519 | 4.5 | 4.01 | 0 | 54 | 2 | -1 | Planum Polare |
| red_v_green |  |  |  |  | 0.981 | 0.877 | 3.52 | 3.26 | 0.001 | 48 | -4 | 5 | Central Opercular Cortex |
| red_v_green | 0.486 | 0.546 | 19 | 0.11 | 0.393 | 0.519 | 4.42 | 3.96 | 0 | 27 | 14 | 38 | Superior Frontal Gyrus |
| red_v_green | 0.038 | 0.087 | 66 | 0.006 | 0.404 | 0.519 | 4.41 | 3.95 | 0 | -24 | 26 | 47 | Superior Frontal Gyrus |
| red_v_green |  |  |  |  | 0.704 | 0.549 | 4.05 | 3.68 | 0 | -15 | 35 | 41 | Superior Frontal Gyrus |
| red_v_green | 0.001 | 0.004 | 143 | 0 | 0.503 | 0.549 | 4.28 | 3.85 | 0 | -51 | -1 | 2 | Central Opercular Cortex |
| red_v_green |  |  |  |  | 0.515 | 0.549 | 4.27 | 3.84 | 0 | -39 | -13 | -10 | Planum Polare |
| red_v_green |  |  |  |  | 0.529 | 0.549 | 4.25 | 3.83 | 0 | -54 | -16 | 11 | Central Opercular Cortex |
| red_v_green | 0.209 | 0.303 | 34 | 0.039 | 0.535 | 0.549 | 4.25 | 3.83 | 0 | -30 | -31 | 44 | Postcentral Gyrus |
| red_v_green |  |  |  |  | 0.763 | 0.574 | 3.98 | 3.62 | 0 | -39 | -28 | 53 | Postcentral Gyrus |
| red_v_green | 0.234 | 0.303 | 32 | 0.044 | 0.572 | 0.549 | 4.2 | 3.79 | 0 | 39 | -19 | -1 | Insular Cortex |
| red_v_green |  |  |  |  | 0.953 | 0.809 | 3.64 | 3.36 | 0 | 42 | -31 | -7 | Middle Temporal Gyrus |
| red_v_green | 0.262 | 0.307 | 30 | 0.05 | 0.604 | 0.549 | 4.17 | 3.77 | 0 | -33 | -19 | 44 | Precentral Gyrus |
| red_v_green |  |  |  |  | 0.835 | 0.583 | 3.88 | 3.55 | 0 | -42 | -25 | 41 | Postcentral Gyrus |
| red_v_gold | 0.024 | 0.049 | 75 | 0.004 | 0.151 | 0.37 | 4.86 | 4.27 | 0 | 12 | -97 | -1 | Occipital Pole |
| red_v_gold |  |  |  |  | 0.244 | 0.37 | 4.65 | 4.13 | 0 | 21 | -88 | -10 | Occipital Fusiform Gyrus |
| red_v_gold |  |  |  |  | 0.977 | 0.794 | 3.54 | 3.28 | 0.001 | 12 | -79 | -7 | Lingual Gyrus |
| red_v_gold | 0.57 | 0.719 | 16 | 0.14 | 0.861 | 0.794 | 3.84 | 3.51 | 0 | -9 | -94 | -1 | Occipital Pole |
| red_v_mtx | 0.293 | 0.388 | 28 | 0.057 | 0.112 | 0.642 | 4.99 | 4.36 | 0 | 39 | -85 | -4 | Lateral Occipital Cortex, inferior division |
| red_v_mtx |  |  |  |  | 0.796 | 0.871 | 3.94 | 3.59 | 0 | 21 | -97 | -4 | Occipital Pole |
| red_v_mtx |  |  |  |  | 0.842 | 0.893 | 3.87 | 3.54 | 0 | 27 | 26 | -10 | Frontal Orbital Cortex |
| red_v_mtx | 0.004 | 0.019 | 113 | 0.001 | 0.157 | 0.642 | 4.85 | 4.26 | 0 | 24 | 56 | 23 | Frontal Pole |
| red_v_mtx |  |  |  |  | 0.382 | 0.642 | 4.44 | 3.97 | 0 | 24 | 62 | 17 | Frontal Pole |
| red_v_mtx |  |  |  |  | 0.767 | 0.871 | 3.97 | 3.62 | 0 | 42 | 47 | 5 | Frontal Pole |
| red_v_mtx | 0.012 | 0.037 | 89 | 0.002 | 0.317 | 0.642 | 4.53 | 4.04 | 0 | 9 | -67 | -7 | Lingual Gyrus |
| red_v_mtx |  |  |  |  | 0.448 | 0.642 | 4.35 | 3.9 | 0 | 24 | -70 | -4 | Occipital Fusiform Gyrus |
| red_v_mtx |  |  |  |  | 0.475 | 0.642 | 4.32 | 3.88 | 0 | 6 | -79 | -7 | Lingual Gyrus |
| red_v_mtx | 0.003 | 0.019 | 124 | 0 | 0.478 | 0.642 | 4.31 | 3.88 | 0 | -39 | -25 | 53 | Postcentral Gyrus |
| red_v_mtx |  |  |  |  | 0.809 | 0.871 | 3.92 | 3.58 | 0 | -27 | -13 | 59 | Precentral Gyrus |
| red_v_mtx |  |  |  |  | 0.967 | 0.97 | 3.59 | 3.32 | 0 | -45 | -28 | 44 | Postcentral Gyrus |
| red_v_mtx | 0.262 | 0.387 | 30 | 0.05 | 0.496 | 0.642 | 4.29 | 3.86 | 0 | 3 | -28 | 23 | Cingulate Gyrus, posterior division |
| red_v_mtx |  |  |  |  | 0.747 | 0.871 | 4 | 3.64 | 0 | -6 | -19 | 29 | Cingulate Gyrus, posterior division |
| red_v_mtx | 0.157 | 0.306 | 39 | 0.028 | 0.497 | 0.642 | 4.29 | 3.86 | 0 | -6 | -34 | 44 | Cingulate Gyrus, posterior division |
| red_v_mtx |  |  |  |  | 0.555 | 0.705 | 4.22 | 3.81 | 0 | -9 | -37 | 32 | Cingulate Gyrus, posterior division |
| red_v_mtx | 0.06 | 0.137 | 57 | 0.01 | 0.747 | 0.871 | 4 | 3.64 | 0 | 39 | -82 | 14 | Lateral Occipital Cortex, superior division |
| red_v_mtx |  |  |  |  | 0.794 | 0.871 | 3.94 | 3.59 | 0 | 39 | -70 | 14 | Lateral Occipital Cortex, inferior division |
| red_v_mtx |  |  |  |  | 0.962 | 0.97 | 3.61 | 3.33 | 0 | 48 | -73 | 14 | Lateral Occipital Cortex, inferior division |
| red_v_mtx | 0.601 | 0.734 | 15 | 0.152 | 0.823 | 0.871 | 3.9 | 3.56 | 0 | 12 | 35 | 14 | Cingulate Gyrus, anterior division |
